# Using Unassigned NMR Chemical Shifts to Model RNA Secondary Structure

**DOI:** 10.1101/2022.01.30.478380

**Authors:** Neel Moudgal, Grace Arhin, Aaron T. Frank

## Abstract

NMR-derived chemical shifts are sensitive probes of RNA structure. However, the need to assign NMR spectra hampers their utility as a direct source of structural information. In this report, we describe a simple method that uses unassigned 2D NMR spectra to model the secondary structure of RNAs. Similar to assigned chemical shifts, we could use unassigned chemical shift data to reweight conformational libraries such that the highest weighted structure closely resembles their reference NMR structure. Furthermore, the application of our approach to the 3’- and 5’-UTR of the SARS-CoV-2 genome yields structures that are, for the most part, consistent with the secondary structure models derived from chemical probing data. Therefore, we expect the framework we describe here will be useful as a general strategy for rapidly generating preliminary structural RNA models directly from unassigned 2D NMR spectra. As we demonstrated for the 337-nt and 472-nt UTRs of SARS-CoV-2, our approach could be especially valuable for modeling the secondary structures of large RNA.

## Introduction

Advancements in our understanding of molecular biology highlight that RNAs play critical cellular roles that go far beyond simply acting as passive information carriers. The ability of some RNA to change structure is often the basis for them executing complex biological function(s).^1,2^ Therefore, determining RNA structure is crucial to building up an intuition about relationships between their sequence, structure, and function. Determining RNA structure begins with their sequence, then their secondary structure(s), that is, the set of canonical and non-canonical base pairs that these sequences form, and then the specific three-dimensional (3D) atomic structure(s) they populate. Thus, predicting the secondary structure of RNA from their sequence is the first and crucial step in understanding RNA structure-function relationships.

For fast and accurate secondary structure prediction from RNA sequence, thermodynamic models that predict the secondary structure with the minimum free energy among all possible secondary structures are utilized.^3^ However, the prediction accuracy of existing thermodynamic models degrades as RNA sequences become longer.^4^ Integrating computational methods with sparse experimental data from various sources has provided a new paradigm for low to medium-resolution structural studies.^5^ Within this integrative framework, experimental data can be coupled with structure prediction algorithms to predict structure conditioned on available data. Notably, several techniques have been developed that utilize NMR chemical shifts to guide the modeling of both the secondary (2D)^6^ and atomic (3D) structures of RNA.^7–9^

Building off previous work,^10^ in which we used *assigned* NMR chemical shifts to pre-dict structure, we implemented and tested a conformational reweighting framework to model RNA secondary structure using *unassigned* NMR chemical shift data. Our motivation for this work is that assigning chemical shifts is labor-intensive, which limits the use of assigned chemical shifts as experimental inputs into integrative modeling pipelines. Methods that can directly incorporate unassigned chemical shift information into the modeling process are attractive because they could help bypass tedious peak assignment and kickstart and accelerate structural studies of novel RNAs.^11^

In what follows, we first describe our implementation of a histogram-based reweighting procedure that one could use to guide secondary structure prediction using unassigned chemical shift data. Second, we report the results of several benchmark tests, in which we use unassigned chemical shift data to resolve the native structure within diverse conformational libraries. Finally, we describe our unprecedented use of unassigned data to model the 3’- and 5’-UTR of the SARS-CoV-2 RNA elements. Overall, our results indicate that the reweighting framework we have implemented allows one to use raw unassigned NMR chemical shift data to accurately predict RNA secondary structure.

## Methods

### Histogram-based chemical shifts reweighting

In this study, we used the Bayesian Maximum Entropy (BME) approach^12–15^ to reweight conformational libraries using 2D chemical shift histograms. By using histograms, we completely by-pass the need for chemical shift assignments and simply rely on counts within common histograms as the basis for comparing chemical shifts computed from structures with actual chemical shift data. In what follows, we will refer to this approach as unassigned chemical shift-based BME or **u**CS-BME (Figure 1). In **u**CS-BME, each conformer in a conformational library is initially assumed to equal weights (i.e, priors),

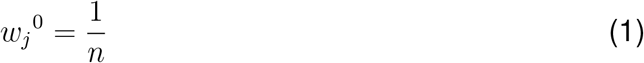

*n* is the total number of conformers in the library and *w*_*j*_^0^ is the initial weight attached to *j*th conformer. During the optimization, conformers are assigned new weights based on how best their back-calculated observables agree with the available experimental observable. In our case, the observables correspond to histogram counts *H*_*i*_(*x*_*j*_) in a specific bin *i*, which depends on the structure *x*_*j*_. The new optimal weights (*w*_*j*_^*\**^) can be expressed as

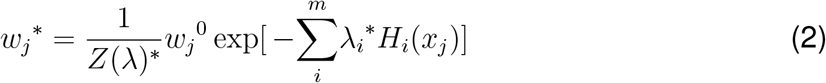

and they are the new assigned weights that minimize the cost function that quantifies the agreement between histogram counts in simulated and experimental 2D spectra. From previous studies,^16–18^ it can be shown that the optimal weights will minimize the following loss function:

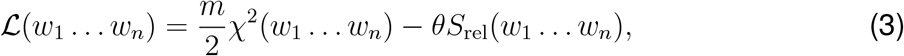

in which:

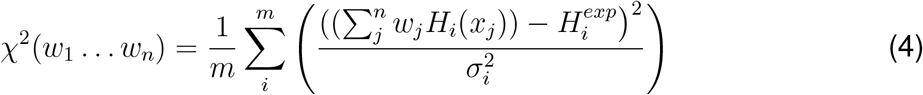

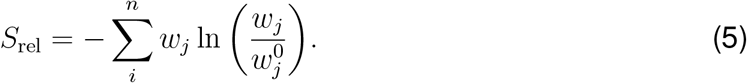

**Figure 1:**
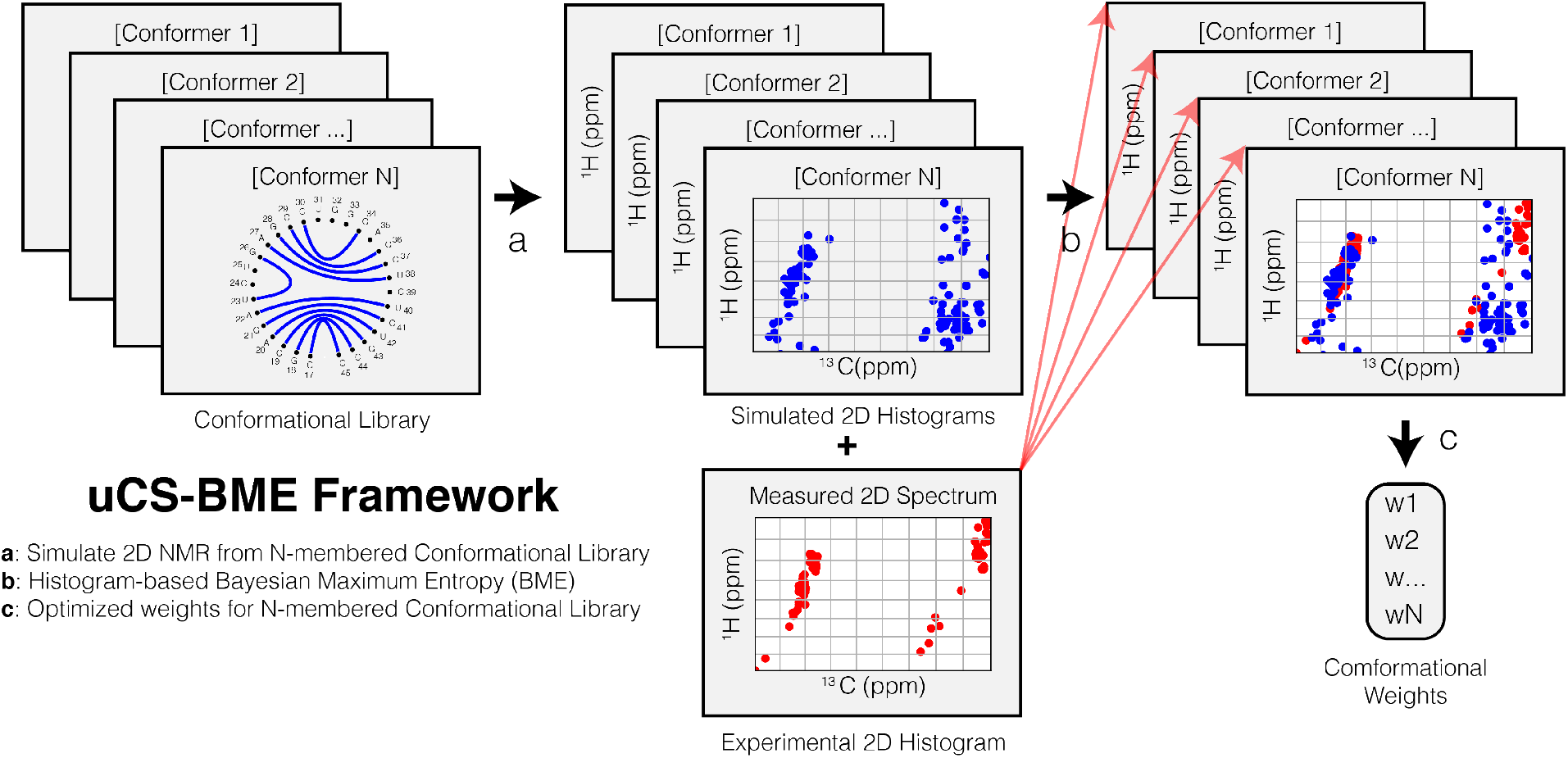
Overview of the uCS-BME framework. (a) We compute chemical shifts for each conformer in an *N* -membered conformational library and then generate a (simulated) 2D chemical shift histogram. (b) We then use the collection of *N* simulated histograms and the corresponding experimental histogram as inputs into the Bayesian Maximum Entropy (BME) reweighting technique, where the goal is to reweight peak counts within individual bins of the simulated 2D histogram so that they agree with peak counts within individual bins of the experimentally-derived 2D histogram. (c) After BME optimization, we extract the optimal weights and define them as the conformation weights in the ensemble.

Here, 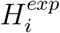 and *H*_*i*_(*x*_*j*_) are the histogram count in bin *i* in a 2D histogram derived from experiments and back-calculated from *j*^th^ conformer in a library, respectively. *σ*_*i*_ is the uncertainty in the histogram count in bin *i*. By setting the bin size in each dimension of the 2D histogram to be a constant that is equal to the expected error in computed chemical shifts, we could then set *σ*_*i*_ to 1, provided that uncertainties in computed chemical shifts are much larger than those in experimentally measured chemical shifts. In the loss function ℒ (Eq. 3), the first term (*χ*^2^) describes the agreement between the experimental data and the back-calculated properties from structure models and the second term (*S*_rel_) describes the deviation of the new weights from the original weights. By optimizing Eq. 3, we could identify weights *w*_*j*_^*\**^ that deviate from *w*_*j*_^0^ only to the point that can minimize *χ*^2^ to within expected errors. The extent to which *w*_*j*_^*\**^ deviate from *w*_*j*_^0^ during the optimization is controlled by *θ*, with increasing values minimizing the deviation and small values allowing larger deviations. *θ* is not known *a priori* and so can be viewed as a hyperparameter whose value optimal must be determined. In this study, we use a cross-validation approach to determine the optimal value of *θ* (see below). To carry out this histogram-based reweighting we utilized the BME code (https://github.com/KULL-Centre/BME) of Bottaro et al^15^.

### Benchmarking histogram-based chemical shifts reweighting

#### Conformational library

To test the ability of unassigned chemical shift-based reweighting to resolve RNA secondary structures, we applied histogram-based BME to reweight the conformational libraries of 12 RNAs based on their sets of measured and computed chemical shifts. The conformational libraries contained sub-optimal conformers generated from RNA sequence using the tool *MC-Fold* from the MC-Sym suite.^19^ Using *MC-Fold*, we generated ten different structures whose folding free energies are within 30% of the lowest energy structure. The exception was the fluoride riboswitch for which we combined decoys generated from *AllSub* from the RNAstructure modeling suite and *MC-Fold* to ensure the library of conformers was diverse, containing both pseudoknotted and non-pseudoknotted structures. The structures were then combined with the native structure to form the final conformational library. In cases where the native, reference NMR structure contained non-canonical base pairs, then both structures with and without the non-canonical base pairs are included the final conformational library.

#### Chemical shift calculation

Next, for each RNA, secondary structure to chemical shift (SS2CS) predictors was used to predict the chemical shifts of all non-exchangeable ^1^H and ^13^C nuclei in each conformer within the conformational libraries.^10^ The computed ^1^H and ^13^C chemical shifts were then used to generate a 2D chemical shift histogram, with ^1^H in the first dimension and ^13^C in the second dimension. A 2D histogram using the experimental ^1^H and ^13^C chemical shifts was also generated. In both cases, the bin size in the ^1^H and ^13^C was set to 0.17 and 0.86 ppm, respectively. These values correspond to the expected (average) accuracy with which chemical shifts for ^1^H and ^13^C nuclei are computed by the SS2CS predictors.^10^ After generating both sets of histograms, we then applied histogram-based BME to assign weights to individual conformers in the conformational libraries.

#### Choosing the optimal value of *θ*

According to Eq. 3, *θ* is a global scaling factor that controls the relative contribution of the entropy terms in the overall loss function *L*. It reflects the trade-off between two terms: (1) *χ*^2^, which is the agreement between experimental data and predicted chemical shifts; (2) *S*_rel_, which is the deviation of the new distribution from the original uniform distribution. The smaller *θ* is, the more *L* is dependent on *χ*^2^. The optimal *θ* was chosen using a five-fold cross-validation approach using the theta_scan functionality in the Bottaro et al.’s BME code (https://github.com/KULL-Centre/BME). Briefly, for each *θ* value between 1.0 and 200.0, we randomly divided the experimental and simulated datasets into a training set and a testing set. After performing BME optimization on the training set using that *θ* value, we generated a set of optimal weights and calculated the value of *χ*^2^ for the testing set given that set of weights. This process was repeated five times, each time randomly choosing a new training and testing set. The optimal *θ* was chosen as the one that minimized average testing error, here the errors in histogram counts.

## Results and Discussion

### uCS-BME weights are similar to CS-BME weights

We recently showed that one could reweight conformational libraries of RNA structures using assigned chemical shift (CS) data and the Bayesian/Maximum Entropy (CS-BME) technique. We found that the highest weighted structure in the CS-BME ensemble closely resembles the reference NMR structure.^10^ To explore whether one can apply the BME framework using unassigned chemical shift data, we implemented a uCS-BME framework. Instead of assigned chemical shift data, uCS-BME uses unassigned chemical shifts (uCS) to reweight 2D conformational libraries using BME.

To accomplish this, we first mimicked unassigned 2D NMR spectra by generating ^1^H/^13^C 2D histograms from assigned measured and computed chemical shift data. For this we paired H1’, H2’, H3’, H4’, H5’, H5”, H2, H5, H6, and H8 with their corresponding carbon peaks, namely, C1’, C2’, C3’, C4’, C5’, C2, C5, C6, and C8. We then generated histograms by binning the H and C dimension. Next, we set the bin width for the H and C dimension to be equal to 0.17 ppm and 0.86 ppm, respectively, which is the inherent accuracy with which one could compute H and C chemical shifts from 2D structures of RNA using a set recently SS2CS models.^10^ We then carried out BME by comparing 2D histograms generated from measured chemical shifts with those computed from the structures in conformational libraries associated with 12 benchmark RNAs. In each case, we used 5-fold cross-validation to identify the optimal *θ*, a hyperparameter contained within the BME optimization function (Eq. 3; Figure S1-S12).

With the optimal *θ*, Δ*χ*^2^, the difference between *χ*^2^ after and before uCS-BME, were all negative, with an average of -0.804 (Table 1). The negative Δ*χ*^2^ indicated that uCS-BME resulted in weights that allowed us to better explain the available data. That is, uCS-BME resulted in reweighted histograms that more closely resembled those derived from measured chemical shifts than the prior model in which each structure in the conformational library was equally weighted. Shown in Figure 2 are bar plots comparing the weights computed using CS-BME and uCS-BME. Across the conformational libraries, the weights derived using CS-BME and uCS-BME were similar, exhibiting high Kendall *τ* ranking coefficients that ranged between 0.59 and 1.00, with a mean of 0.88 (Figure 2 and Table 1).

**Table 1:** Summary of uCS-BME ensembles of RNAs in the validation set using unassigned NMR data. For comparison, results obtained using CS-BME are included in parentheses.

| PDB ID | Size | $N$ | $\theta$ | $\Delta\chi^2$ | $\tau$ | $\phi$ | Max. Weight | TPR (%) | PPV (%) |
| --- | --- | --- | --- | --- | --- | --- | --- | --- | --- |
| 1HWQ | 30 | 11 | 1.00 (4.99) | -1.64 (-1.64) | 0.93 | 0.09 (0.09) | 1.00 (1.00) | 75.00 (75.00) | 100.00 (100.00) |
| 1YMO | 47 | 12 | 12.47 (1.00) | -1.57 (-1.58) | 1.00 | 0.14 (0.12) | 0.82 (0.89) | 93.75 (93.75) | 100.00 (100.00) |
| 2L3E | 35 | 11 | 1.00 (1.00) | -0.71 (-0.71) | 1.00 | 0.10 (0.10) | 0.98 (0.98) | 85.71 (85.71) | 100.00 (100.00) |
| 2LU0 | 49 | 12 | 200.00 (1.00) | -1.05 (-1.17) | 0.70 | 0.34 (0.13) | 0.62 (0.84) | 84.21 (84.21) | 100.00 (100.00) |
| 2N6X | 43 | 12 | 1.00 (1.00) | -1.30 (-1.30) | 1.00 | 0.11 (0.11) | 0.94 (0.94) | 93.75 (93.75) | 100.00 (100.00) |
| 2N7X | 23 | 11 | 16.96 (1.00) | -0.33 (-0.37) | 0.85 | 0.42 (0.18) | 0.42 (0.62) | 100.00 (100.00) | 100.00 (100.00) |
| 2N82 | 23 | 12 | 200.00 (1.00) | -0.36 (-0.85) | 0.70 | 0.90 (0.19) | 0.19 (0.71) | 66.67 (66.67) | 100.00 (100.00) |
| 2NBY | 39 | 11 | 2.50 (1.00) | -0.57 (-0.57) | 0.93 | 0.12 (0.10) | 0.95 (0.99) | 88.24 (88.24) | 100.00 (100.00) |
| 2NC0 | 28 | 11 | 1.00 (1.00) | -0.68 (-0.68) | 1.00 | 0.11 (0.11) | 0.97 (0.97) | 83.33 (83.33) | 100.00 (100.00) |
| 5KH8 | 47 | 18 | 199.00 (2.50) | -0.28 (-0.52) | 0.59 | 0.87 (0.21) | 0.14 (0.61) | 94.12 (94.12) | 100.00 (100.00) |
| 5KMZ | 31 | 11 | 200.00 (35.41) | -0.16 (-0.33) | 0.96 | 0.96 (0.75) | 0.16 (0.28) | 33.33 (33.33) | 30.77 (30.77) |
| 6GZK | 48 | 13 | 4.99 (200.00) | -1.00 (-0.80) | 0.86 | 0.26 (0.61) | 0.50 (0.37) | 76.19 (76.19) | 100.00 (100.00) |
| mean | 45 | 13 | 69.99 (20.91) | -0.80 (-0.88) | 0.88 | 0.37 (0.22) | 0.64 (0.77) | 81.19 (81.19) | 94.23 (94.23) |

**Figure 2:**
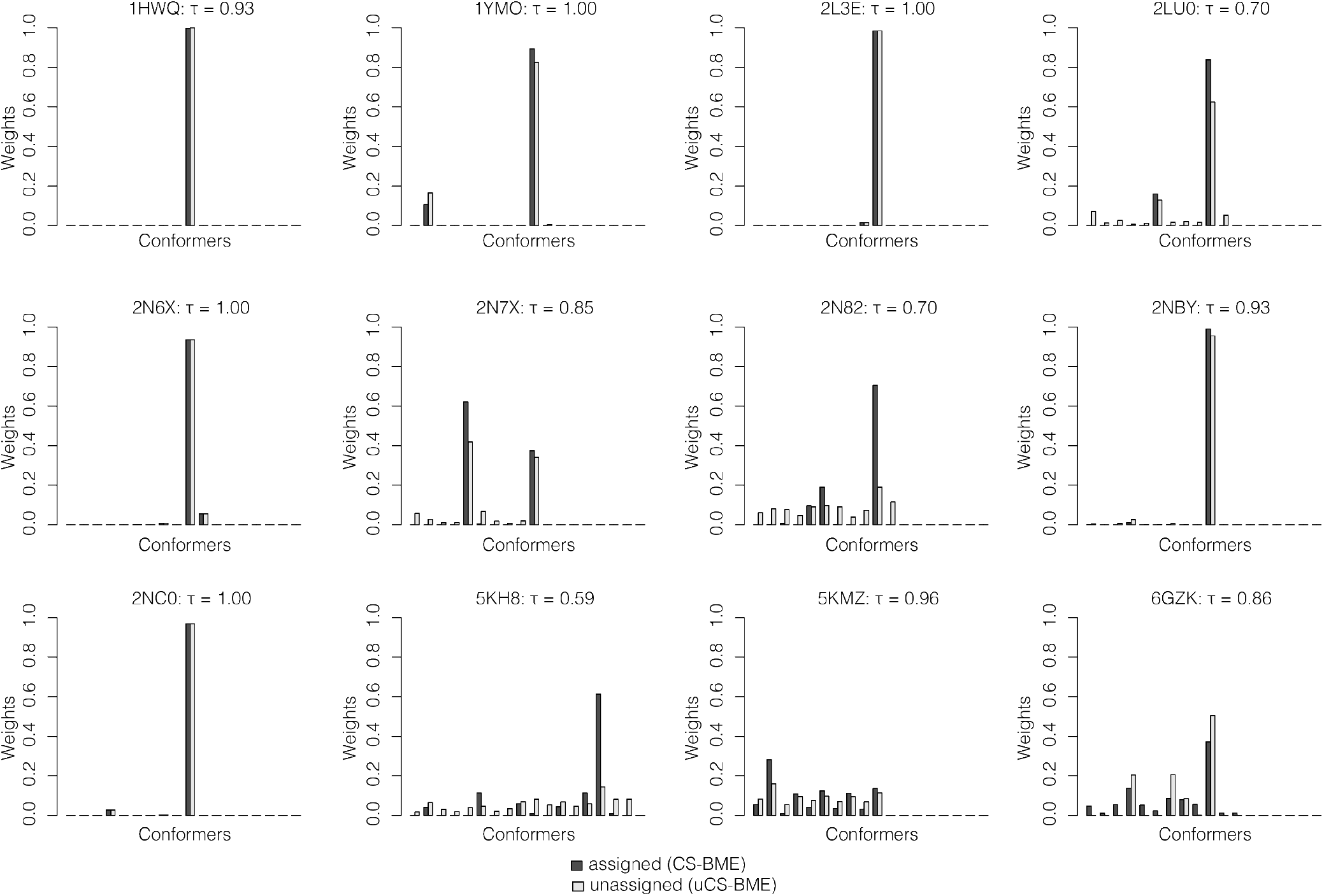
Weight comparison. Barplots comparing weights from CS-BME (assigned) and uCS-BME (unassigned). Individual plots are annotated with the Kendall (*τ*) rank coefficient. *τ* ranking values approaching 1 indicate that the relative ranking of conformers in the library when using assigned weights derived from CS-BME are same as those derived from uCS-BME.

To quantify the distribution of the usCS-BME weights across individual conformational libraries, we computed the effective sample size (*ϕ*).^20,21^ *ϕ* values approaching 0 indicating a single conformer dominate the distribution, whereas values near 1 indicate that the weights are evenly distributed across the conformers in a library. Across the 12 RNAs, uCS-BME *ϕ* ranged between 0.09 and 0.96, with a mean of 0.37 (Table 1). By comparison, the corresponding values using assigned chemical shifts (CS-BME) were 0.09 and 0.75, with a mean of 0.22 (Table 1). Therefore, although the weights from uCS-BME were similar to those derived using CS-BME, uCS-BME-derived weights were more distributed across the set of conformers.

### Like CS-BME, structures with the highest uCS-BME weights closely resemble reference NMR structures

To check whether the highest weighted structures resembled the reference NMR structures, we computed their true positive rate (TPR or sensitivity) and positive predicted value (PPV or precision). The TPR captures the fraction of the “true” base pairs in the reference NMR structure that were found in the highest weighted structure. On the other hand, the PPV captures the fraction of base pairs in the highest weighted structure that are found in the reference NMR structure. In both cases, values near 100.00% indicate that the two structures are highly similar. Across the 12 benchmark RNAs, the highest weighted uCS-BME structures exhibited TPR that ranged between 33.33 and 100.00%, with a mean of 81.19% and PPV that ranged between 30.77 and 100.00%, with a mean of 94.23%, indicating that in general, the highest weighted conformer in the libraries resembled the reference NMR structures (Table 1). (Because the highest weighted uCS-BME structure was the same as the highest weighted CS-BME structure, the TRP and PPV values were identical (Table 1).) These results indicate the highest weighted conformers identified using unassigned data closely resembled the reference NMR structures (Figure S14-S25), and their accuracy was on par with the accuracy achieved using the weights derived from assigned data.

### uCS-BME modeling of the UTRs of SARS-CoV-2 yield structures that are consistent with those derived from DMS footprinting

The results described in the preceding section strongly suggest that the uCS-BME framework could be an effective strategy for using unassigned NMR chemical shifts to guide the structural modeling of RNA. This framework could be particularly advantageous when studying large RNAs that might exhibit significant spectral overlap that hinders unambiguous peak assignment. To explore the utility of uCS-BME in modeling the structure of large RNAs, we next applied uCS-BME to the (337-nt) 3’- and (472-nt) 5’-untranslated regions (UTRs) of the SARS-CoV-2 RNA genome. We were particularly interested in quantifying the extent to which the highest weighted uCS-BME resembled the DMS-derived secondary structure model.^22^ For both the 3’- and 5’-UTRs, we used unassigned imino (G/U) peaks derived from ^1^H, ^1^N TROSY-HSQC (imino) spectrum^22^ to reweight conformational libraries containing the 500 lowest free energy structures. We then computed 2D-histograms representing the 2D NMR spectra were using bin sizes of 1.89 and 0.36 in the 15N and 1H dimensions, respectively. These values corresponded to the mean expected accuracy of (SS2CS) chemical shift predictors that we trained to predict to chemical shifts of N1 and N3 and H1 and H3 nuclei, respectively, from secondary structure models (Table S1).

In Figure 3 we compare the highest uCS-BME weighted structures, and the minimum free energy (MFE) structure the 3’-UTR (Figure 3a and b) and 5’-UTR (3c and d), respectively, to their corresponding DMS-derived secondary structure models. Interestingly, the uCS-BME were on-par (in the case of 3’-UTR) or better agreement (in the case of the 5’-UTR) with DMS-derived secondary structure models than the MFE structures. In the case of the 3’-UTR, the TPR and PPV of the uCS-BME structures relative to the DMS structures were 92.38 and 96.04% (Figure 3a) compared with 91.43 and 95.05% for the MFE structure (Figure 3b). And, in the case of the 5’-UTR, the TPR and PPV of the uCS-BME structures relative to the DMS structures were 88.41 and 77.71% (Figure 3c) compared with 76.06 and 70.47% for the MFE structure (Figure 3d). For the 5’-UTR, the uCS-BME recovers most of the base-pairs in stem-loops (SL) 1 thru 7 (SL1-SL7). The primary disagreement between the uCS-BME and DMS-derived structures occurs within base-pairs associated with residues beyond residue 400, which have previously been predicted to form a pseudoknot.^22^ None of the 500 lower energy structures in the library included pseudoknots. When we exclude these residues, the TPR and PPV of uCS-BME structures relative to the DMS structures increase to 92.19 and 91.47% (Figure 3c) compared with 75.19 and 78.86% for the MFE structure (Figure 3d).

**Figure 3:**
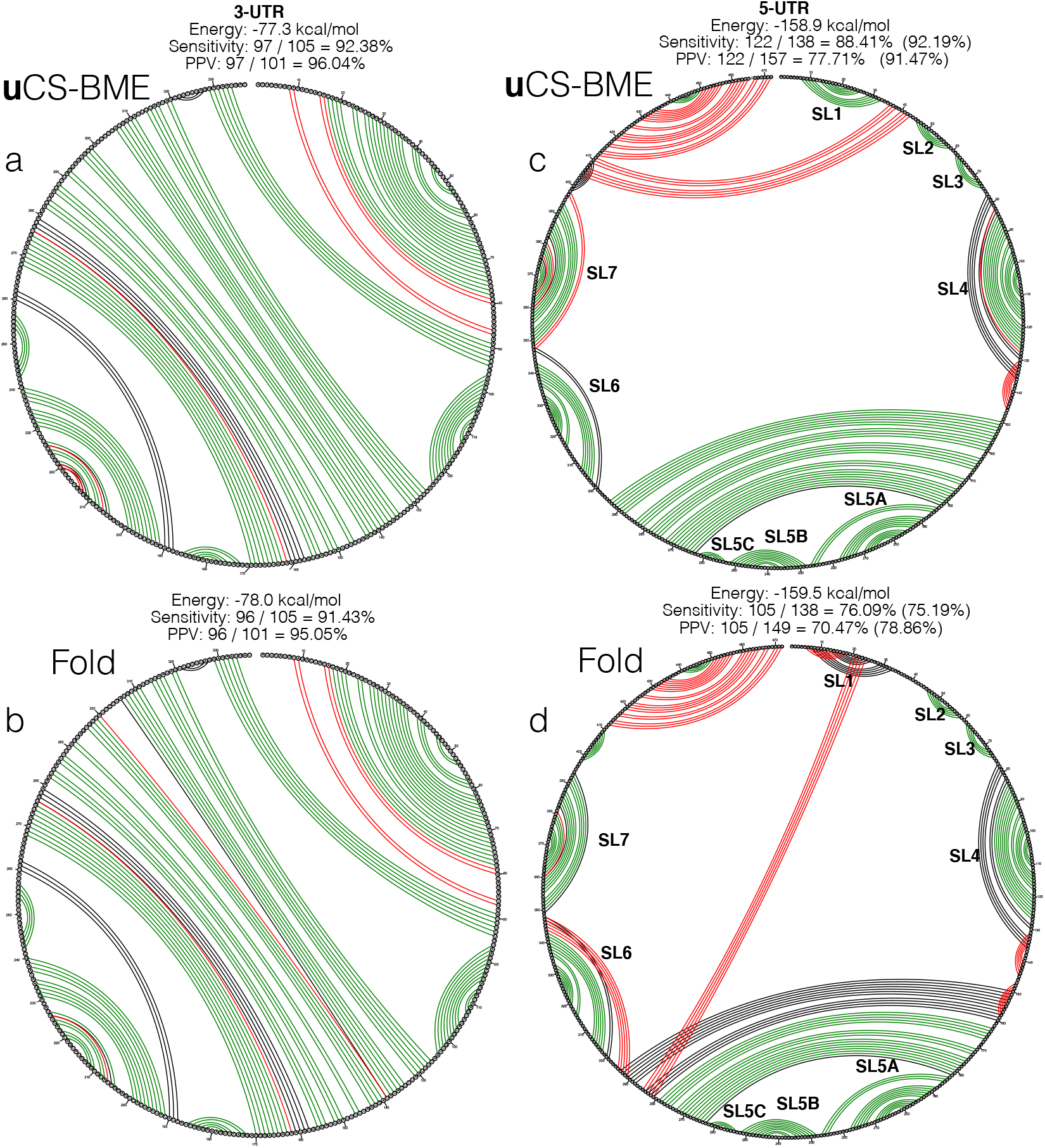
uCS-BME modeling of the 3’-UTR and 5’-UTR of the SARS-CoV-2 RNA genome. Shown are CircleCompare plots for the comparison between DMS-derived secondary structure and (a/c) the highest weighted uCS-BME and (b/d) the MFE structure of the 3’/5’-UTR.

## Discussion and Conclusions

NMR chemical shifts are sensitive probes of structure and dynamics, making them very useful in structural and dynamical studies of flexible biomolecules, like RNA. We recently demonstrated how, using what we referred to as CS-BME, one could integrate assigned NMR chemical shifts with structural prediction tools to determine the 2D ensemble of RNA by Bayesian Maximum Entropy (BME) reweighting.^10^ As important as these chemical shifts are, their assignment is a very labor-intensive process, especially in the case of larger biomolecules where short transverse relaxation (T2) times lead to spectral overlap and line broadening, making it difficult, if not impossible, to unambiguously assign them. In this study, we have introduced uCS-BME reweighting, which facilitates the use of *unassigned* chemical shift data to determine the secondary structure of RNAs. After benchmarking and comparing with CS-BME, we challenged the uCS-BME to predict entire structures of the 5’- and 3’-UTRs of the SARS-COV-2 viral genome. Intriguingly, we found that the highest weighted structure in the ensemble for both the 5’-UTR and 3’-UTR of SARS-COV-2 genome contains many of the base pairs found in the DMS-derived structures of these UTRs; the calculated TPR and PPV for both the 3’-UTR and 5’-UTR structures with respect to the DMS-derived structure is 92.38 and 96.04%, and 88.41 and 77.71% respectively.

Like CS-BME, the performance of this uCS-BME is limited by two main factors. First, the accuracy with which chemical shifts can be computed from structural models, and second, the quality of the initial conformational library being reweighted; that is, the extent to which it contains all of the likely conformational states accessible to a given RNA. As more data become available, it is reasonable to expect that the accuracy with which chemical shifts can be predicted is likely to increase, facilitating more robust comparisons between simulated and measured NMR chemical shifts. Likewise, higher quality conformational libraries will be more accessible as better thermodynamic energy functions are developed as well as better conformational sampling procedures, which allow one to rapidly better sample both non-nested and nested (pseudoknotted) structures.

Furthermore, we acknowledge that inherent in the histogram-based uCS-BME we implemented and tested here are three hyper-parameters, two associated with the bin sizes that are used to generate 2D histograms and the other, *θ*, which appears in the objective function that is optimized during BME reweighting (Eq. 3). The bin sizes we used to generate the 2D histograms were set to the expected accuracy of the chemical shift predictors. For example, for the ^1^H dimension in a histogram, the bin size was set as the mean expected accuracy of the predictors used to estimate ^1^H chemical shifts from secondary structures. These estimates, however, are somewhat biased by the data set over which the performance of the models is evaluated. We found, however, that as it relates to recovering structures that are consistent with known structures, the uCS-BME framework is robust against minor changes in the bin sizes (Table S2-S5). As it relates to choosing *θ*, we employed a 5-fold cross-validation selection approach that is more rigorous than the approach that we previously used for CS-BME (Figure S1-S12).

In spite of these limitations, the results of our study suggest that with more accurate chemical shift predictors and better techniques for sampling complex secondary structures of RNA, frameworks like the uCS-BME we have implemented will provide an avenue for using unassigned NMR chemical shift data to *rapidly* generate structural hypotheses about the likely underlying secondary structure of large RNAs. One could then use such models as inputs to 3D structure prediction pipelines to arrive at an atomic description of the likely structure of these RNAs. Such atomic structures could then be used to rationalize sequence-structure-function relationships of individual RNAs.

## Author Contributions

ATF conceived the project, NM, GA, and ATF conducted the computational experiments; NM, GA, and ATF analyzed the results and wrote the manuscript. All authors reviewed the manuscript.

## Acknowledgments

We thank the members of the Frank lab for many valuable discussions about this work. We would like to express special gratitude to Ziqiao Xu for his enthusiastic support of this project and generous gift of time. We also would like to thank Prof. Dr. Harald Schwalbe, Dr. Christian Richter, and Dr. Anna Wacker for sharing the 2D chemical shift data for the SARS-CoV-2 RNAs. This research was supported by start-up funds from the University of Michigan and by the National Science Foundation grant MCB-2046005 (to ATF).

## Supporting Information

Figure S1, uCS-BME cross-validation plots for 1HWQ; Figure S2, uCS-BME cross-validation plots for 1YMO; Figure S3, uCS-BME cross-validation plots for 2L3E; Figure S4, uCS-BME cross-validation plots for 2LU0; Figure S5, uCS-BME cross-validation plots for 2N6X; Figure S6, uCS-BME cross-validation plots for 2N7X; Figure S7, uCS-BME cross-validation plots for 2N82; Figure S8, uCS-BME cross-validation plots for 2NBY; Figure S9, uCS-BME cross-validation plots for 2NC0; Figure S10, uCS-BME cross-validation plots for 5KH8; Figure S11, uCS-BME cross-validation plots for 5KMZ; Figure S12, uCS-BME cross-validation plots for 6GZK; Figure S25, distribution of differences between usCS-BME and CS-BME-derived weights. Figure S13, CS^2^BP^2^Plots for for 1HWQ; Figure S14, CS^2^BP^2^Plots for for 1YMO; Figure S15, CS^2^BP^2^Plots for for 2L3E; Figure S16, CS^2^BP^2^Plots for for 2LU0; Figure S17, CS^2^BP^2^Plots for for 2N6X; Figure S18, CS^2^BP^2^Plots for for 2N7X; Figure S19, CS^2^BP^2^Plots for for 2N82; Figure S20, CS^2^BP^2^Plots for for 2NBY; Figure S21, CS^2^BP^2^Plots for for 2NC0; Figure S22, CS^2^BP^2^Plots for for 5KH8; Figure S23, CS^2^BP^2^Plots for for 5KMZ; Figure S24, CS^2^BP^2^Plots for for 6GZK; Table S1, accuracy of N1/H1 and N3/H3 SS2CS predictors; Table S2, variations in TPR with histogram bin sizes for the 3’-UTR of SARS-CoV-2; Table S3, variations in PPV with histogram bin sizes for the 3’-UTR of SARS-CoV-2; Table S4, variations in TPR with histogram bin sizes for the 5’-UTR of SARS-CoV-2; Table S5, variations in PPV with histogram bin sizes for the 5’-UTR of SARS-CoV-2.

## TOC IMAGE

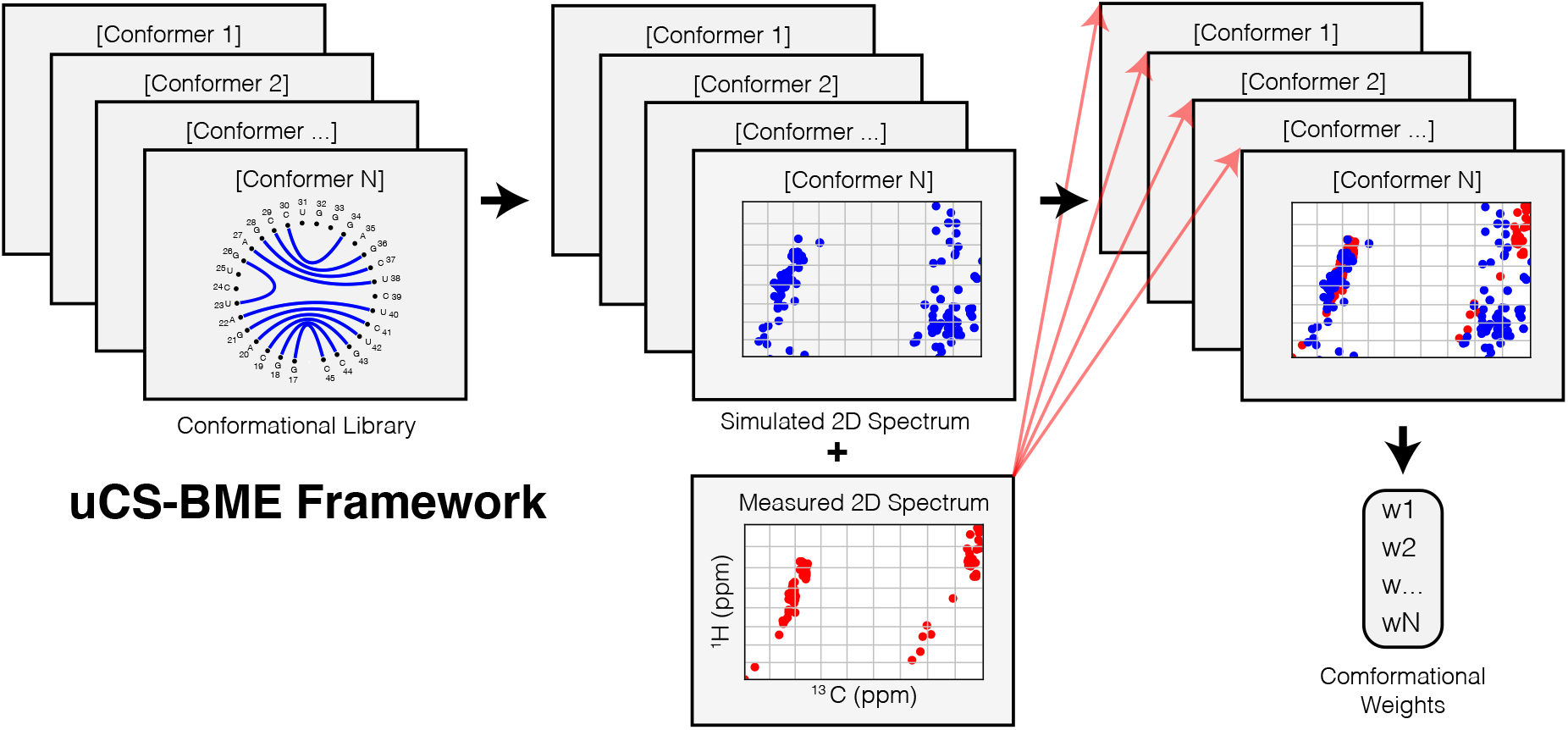

## SUPPORTING INFORMATION

**Figure S1:**
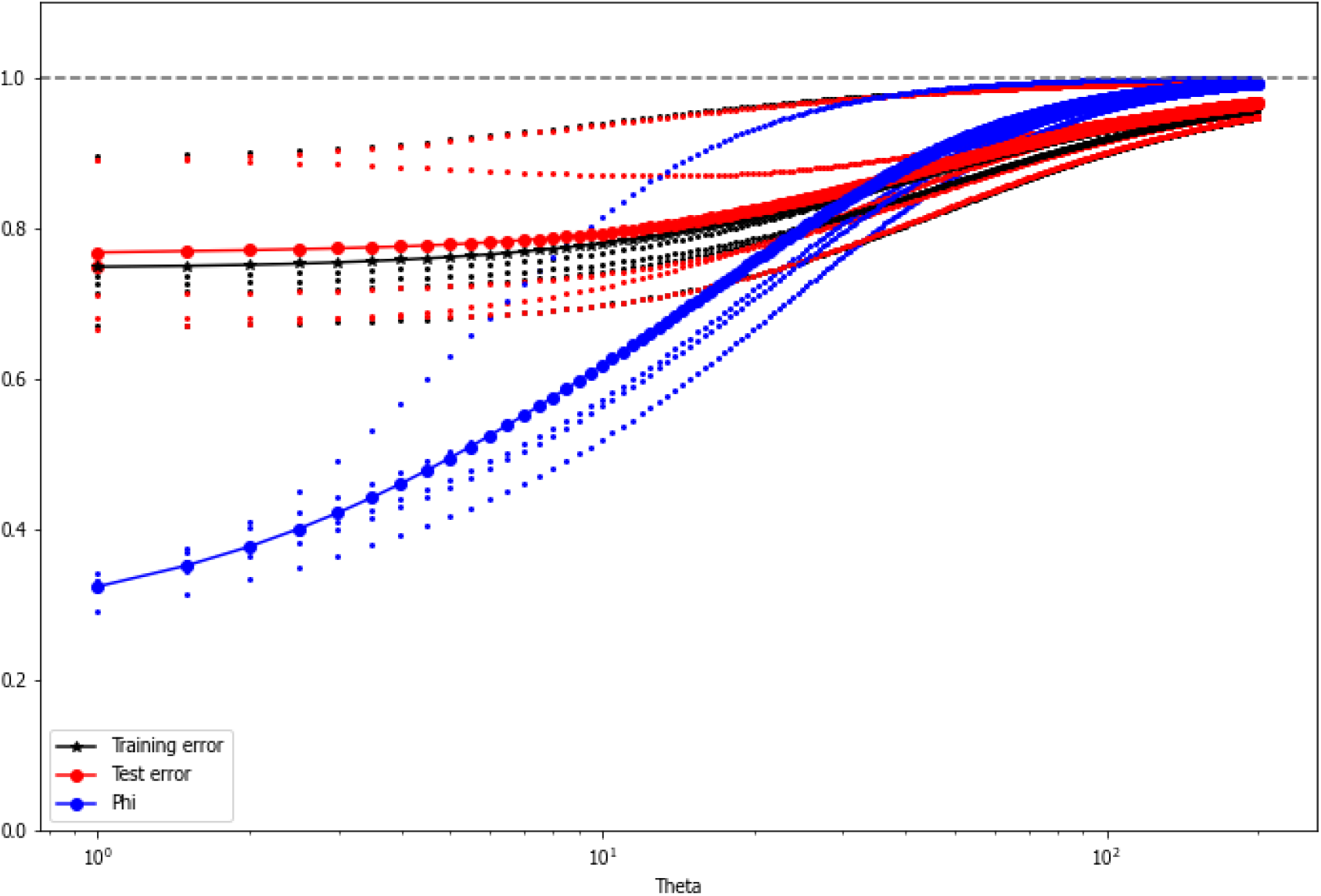
uCS-BME 5-fold cross-validation results that was used to choose the value of *θ* for PDB ID: 1HWQ. The training and testing errors are shown in black and red, respectively. Shown in blue is *ϕ*, the effective ensemble size. The optimal *θ* is chosen as the value that minimizes the average testing error.

**Figure S2:**
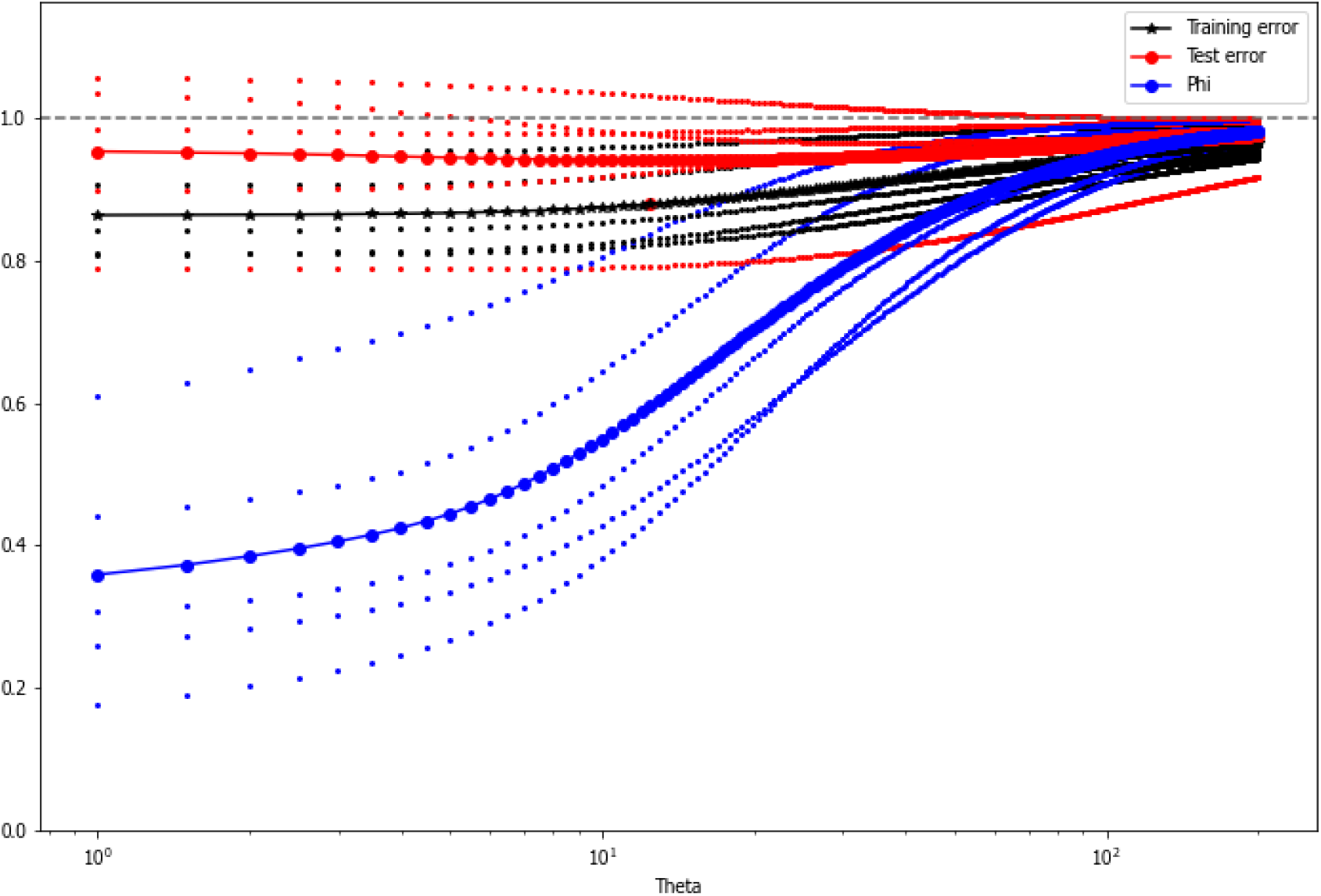
uCS-BME 5-fold cross-validation results that was used to choose the value of *θ* for PDB ID: 1YMO. The training and testing errors are shown in black and red, respectively. Shown in blue is *ϕ*, the effective ensemble size. The optimal *θ* is chosen as the value that minimizes the average testing error.

**Figure S3:**
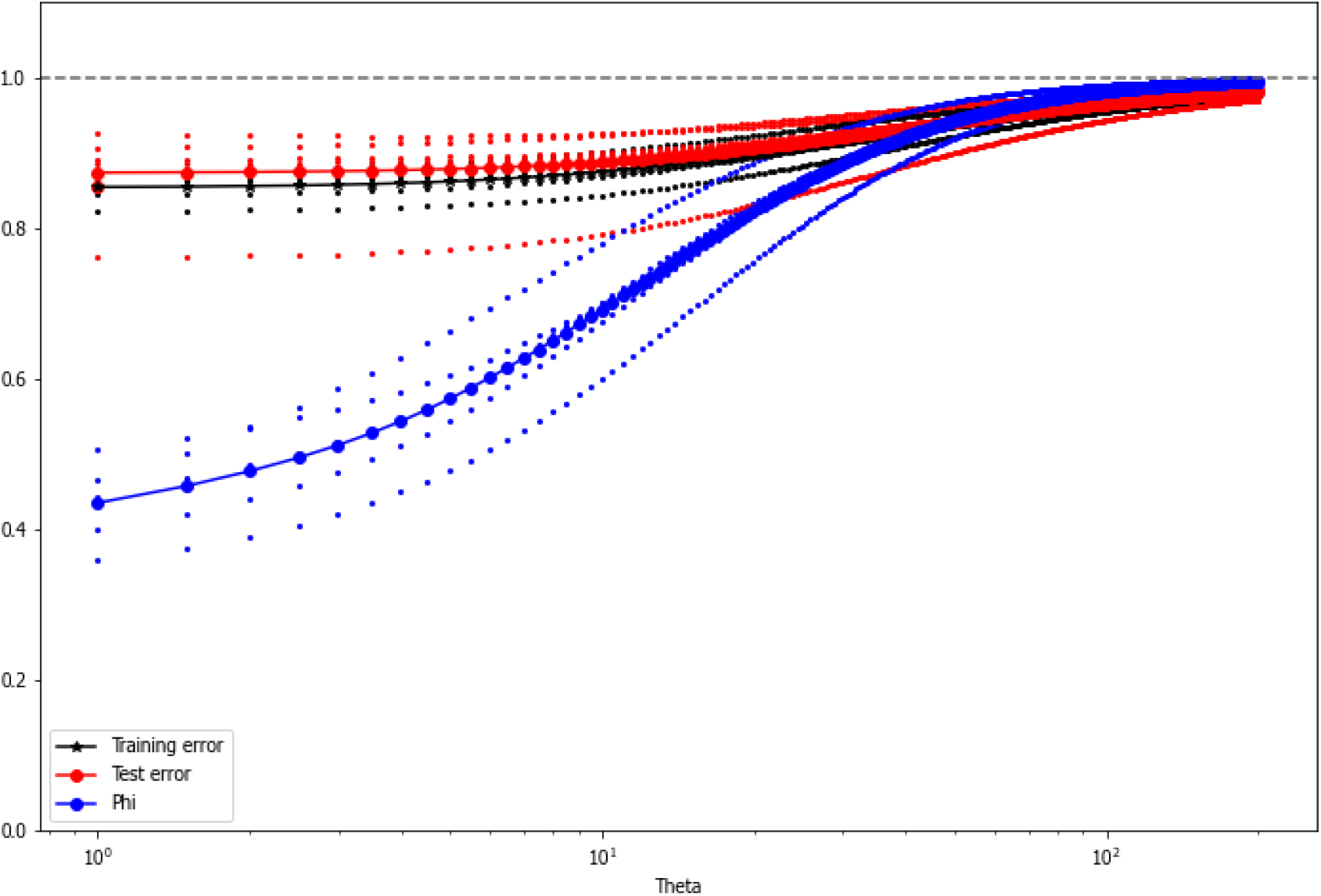
uCS-BME 5-fold cross-validation results that was used to choose the value of *θ* for PDB ID: 2L3E. The training and testing errors are shown in black and red, respectively. Shown in blue is *ϕ*, the effective ensemble size. The optimal *θ* is chosen as the value that minimizes the average testing error.

**Figure S4:**
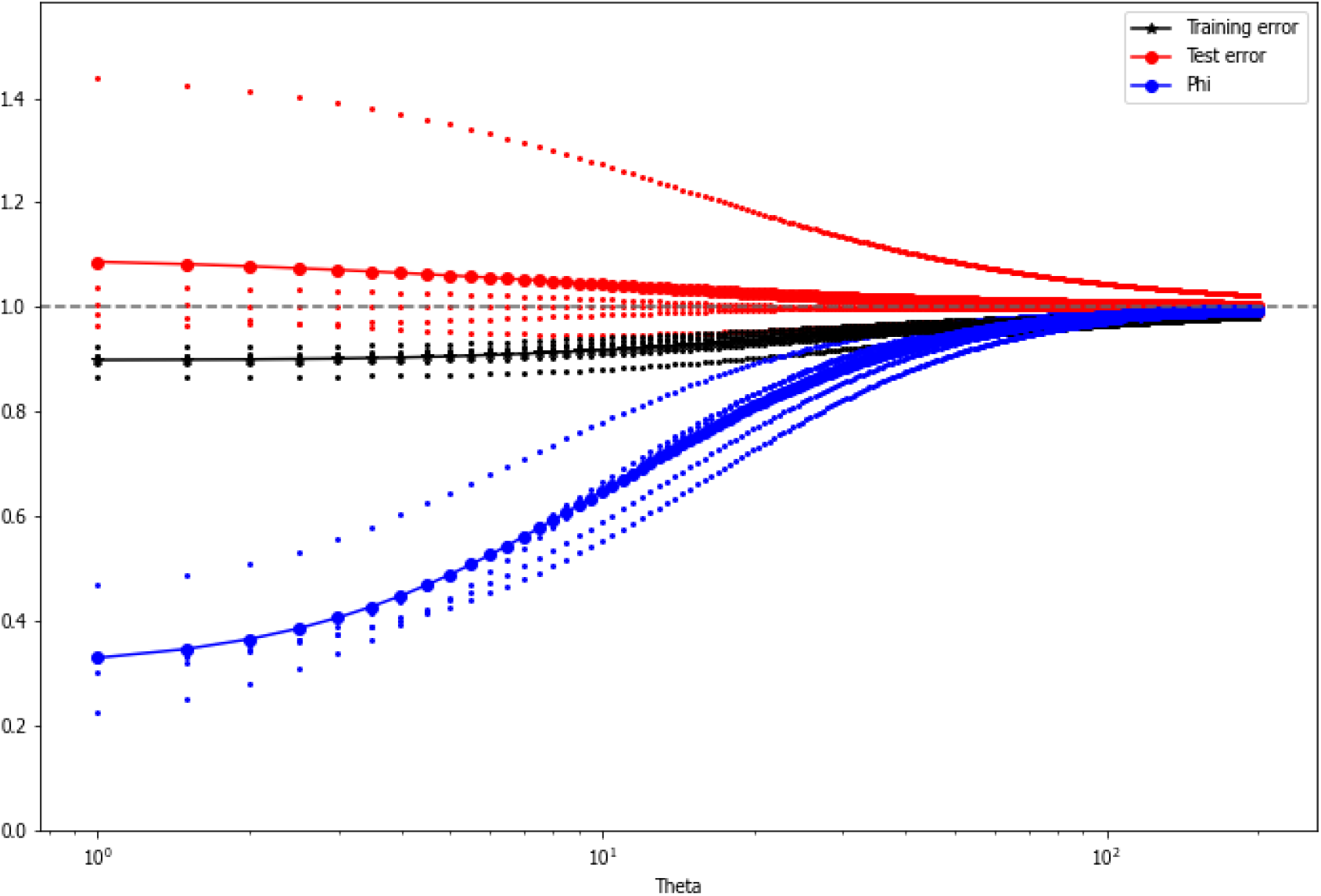
uCS-BME 5-fold cross-validation results that was used to choose the value of *θ* for PDB ID: 2LU0. The training and testing errors are shown in black and red, respectively. Shown in blue is *ϕ*, the effective ensemble size. The optimal *θ* is chosen as the value that minimizes the average testing error.

**Figure S5:**
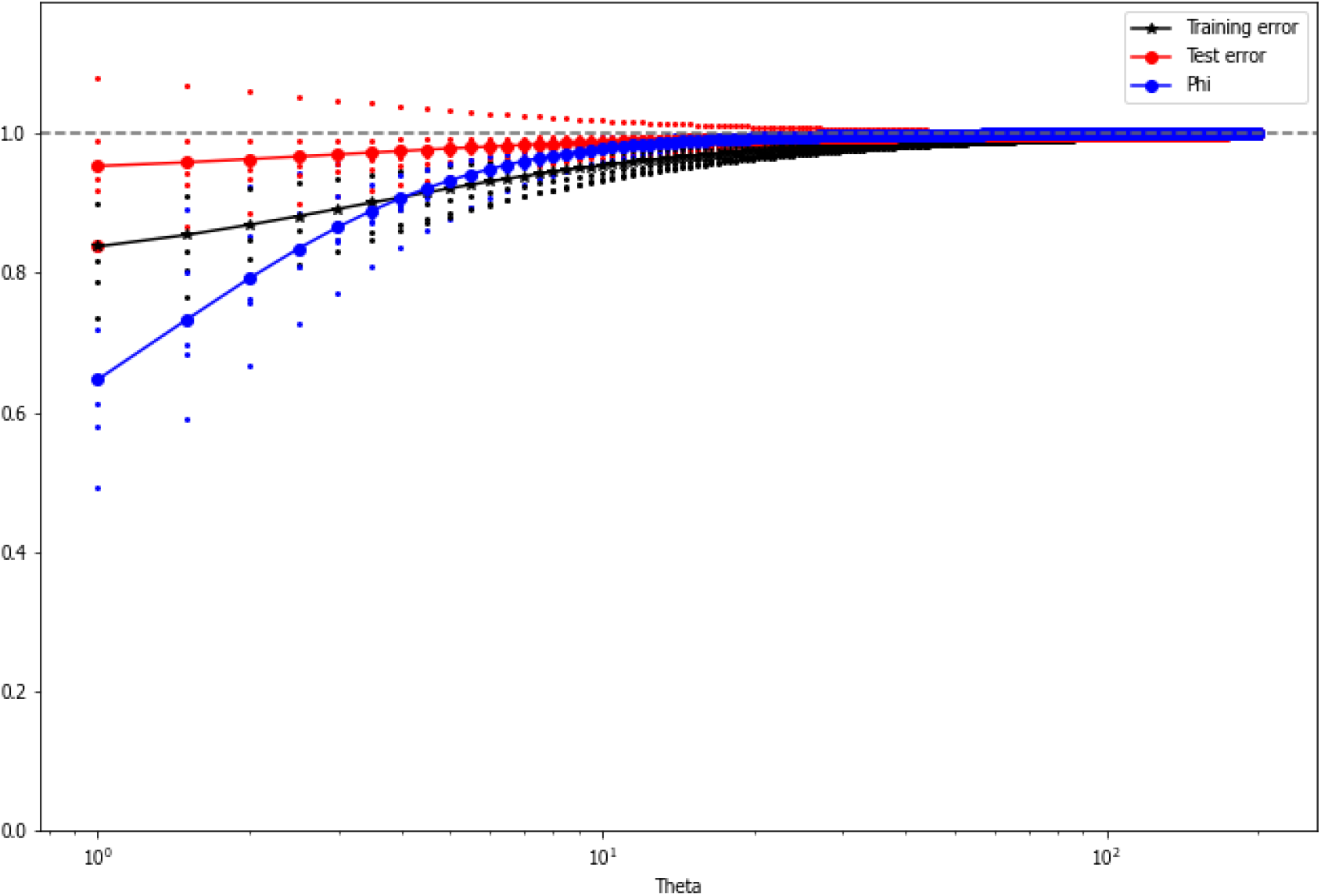
uCS-BME 5-fold cross-validation results that was used to choose the value of *θ* for PDB ID: 2N6X. The training and testing errors are shown in black and red, respectively. Shown in blue is *ϕ*, the effective ensemble size. The optimal *θ* is chosen as the value that minimizes the average testing error.

**Figure S6:**
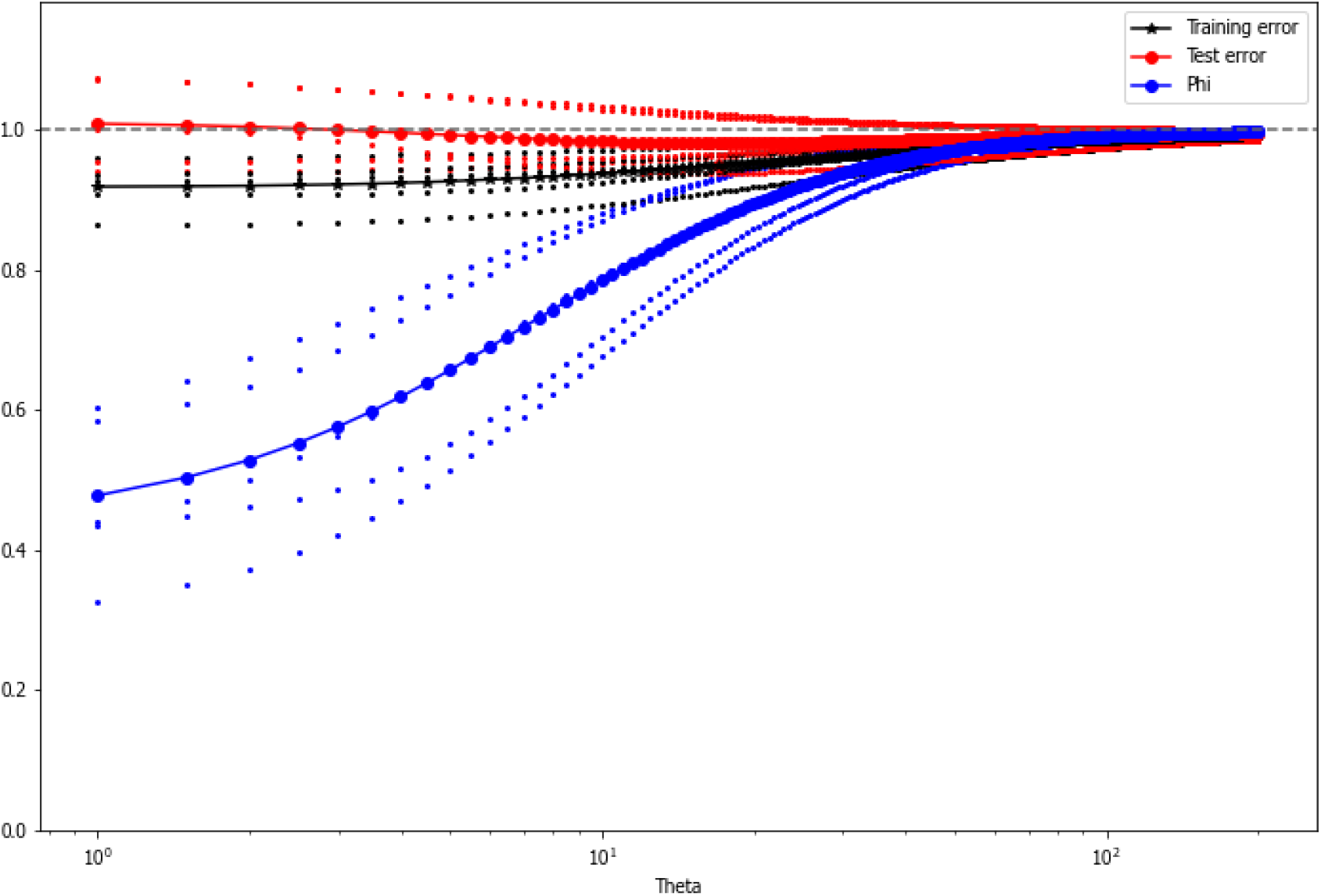
uCS-BME 5-fold cross-validation results that was used to choose the value of *θ* for PDB ID: 2N7X. The training and testing errors are shown in black and red, respectively. Shown in blue is *ϕ*, the effective ensemble size. The optimal *θ* is chosen as the value that minimizes the average testing error.

**Figure S7:**
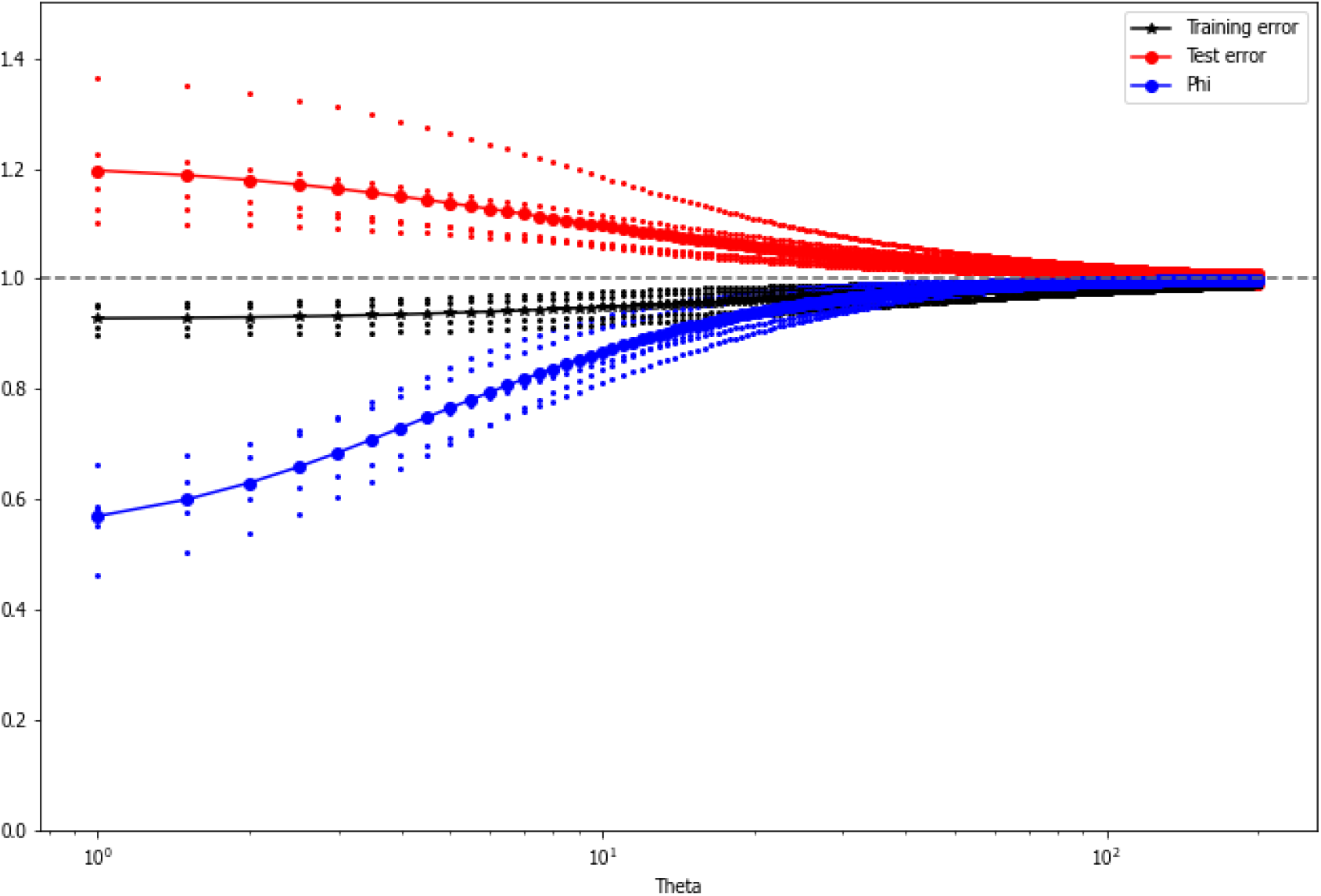
uCS-BME 5-fold cross-validation results that was used to choose the value of *θ* for PDB ID: 2N82. The training and testing errors are shown in black and red, respectively. Shown in blue is *ϕ*, the effective ensemble size. The optimal *θ* is chosen as the value that minimizes the average testing error.

**Figure S8:**
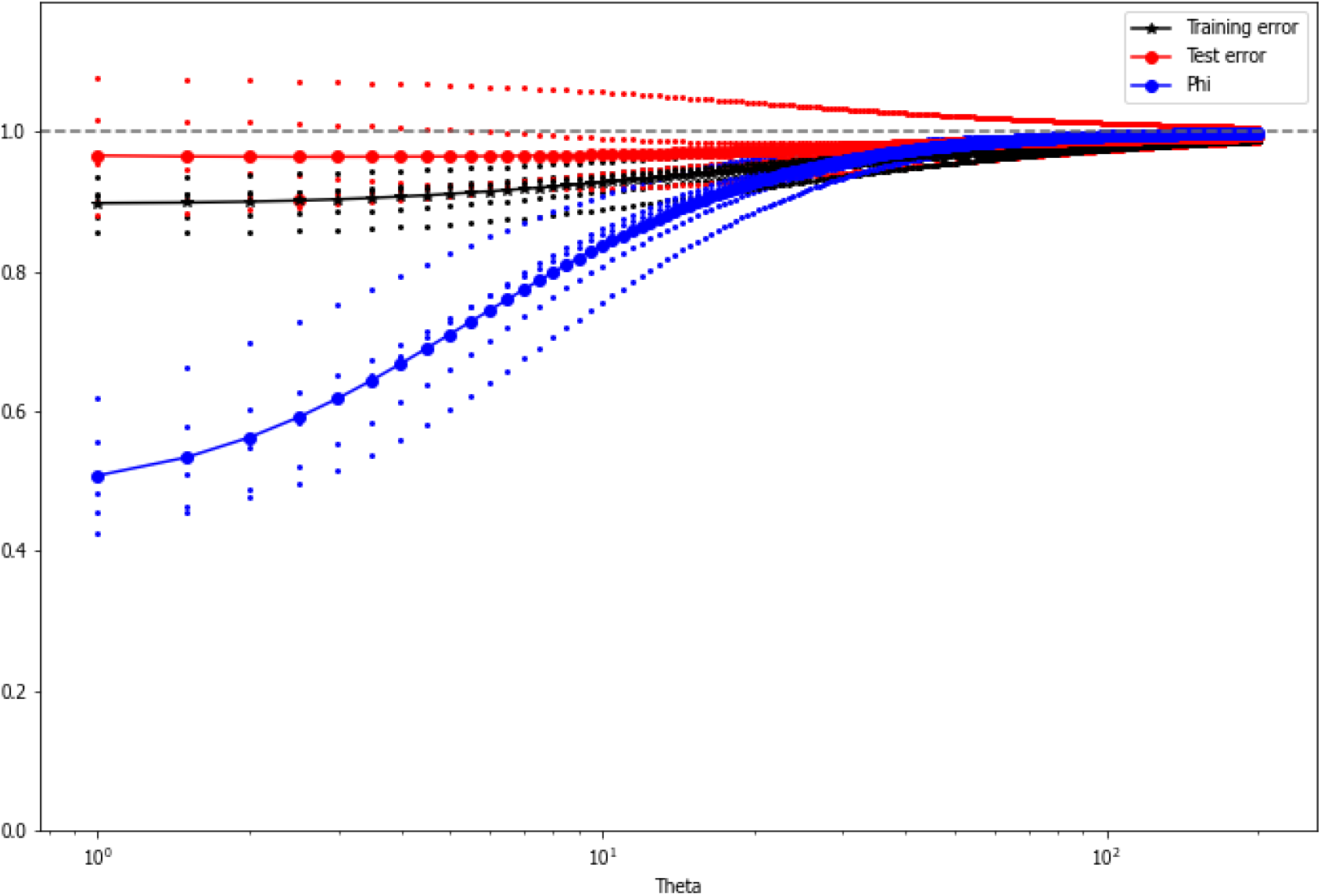
uCS-BME 5-fold cross-validation results that was used to choose the value of *θ* for PDB ID: 2NBY. The training and testing errors are shown in black and red, respectively. Shown in blue is *ϕ*, the effective ensemble size. The optimal *θ* is chosen as the value that minimizes the average testing error.

**Figure S9:**
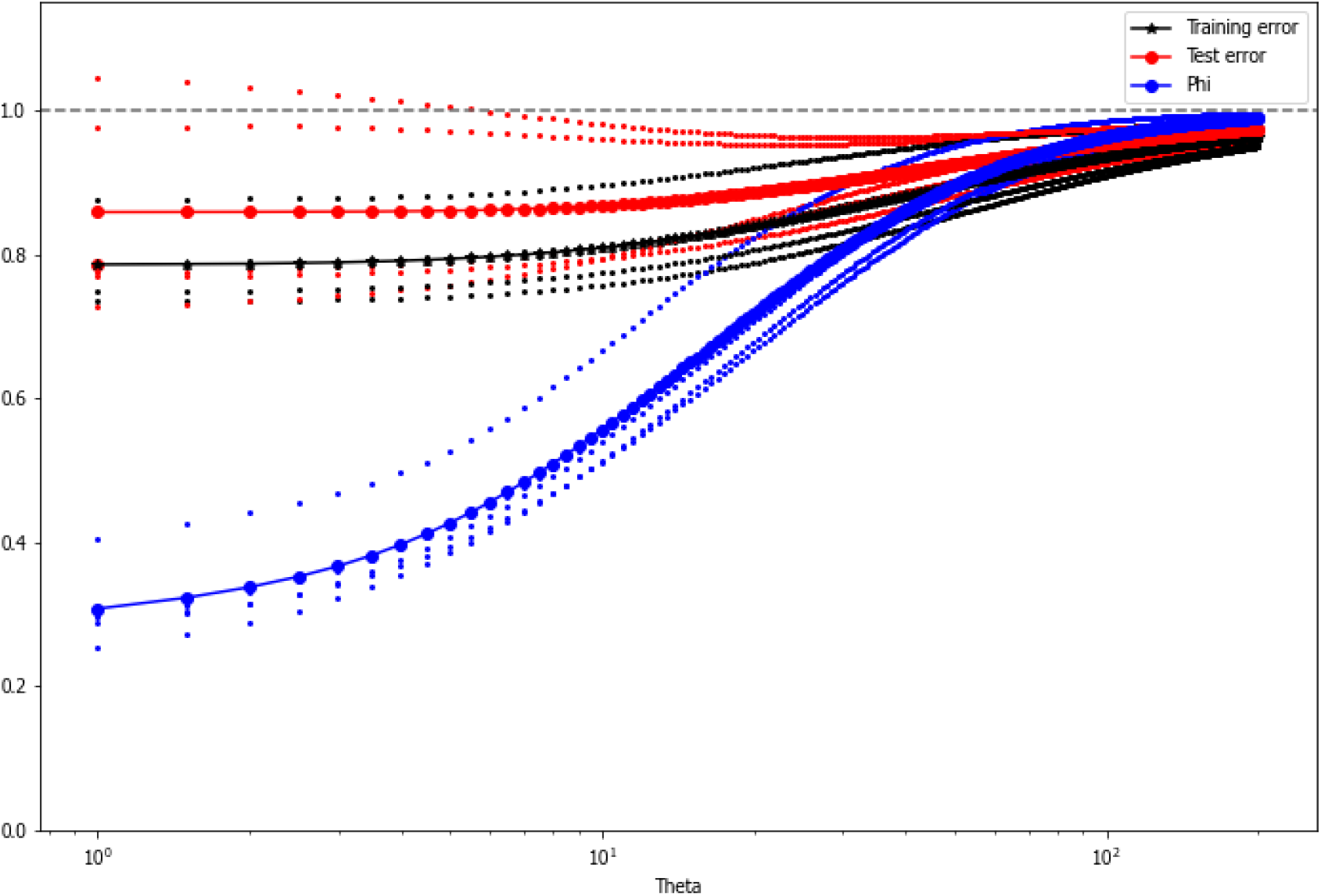
uCS-BME 5-fold cross-validation results that was used to choose the value of *θ* for PDB ID: 2NC0. The training and testing errors are shown in black and red, respectively. Shown in blue is *ϕ*, the effective ensemble size. The optimal *θ* is chosen as the value that minimizes the average testing error.

**Figure S10:**
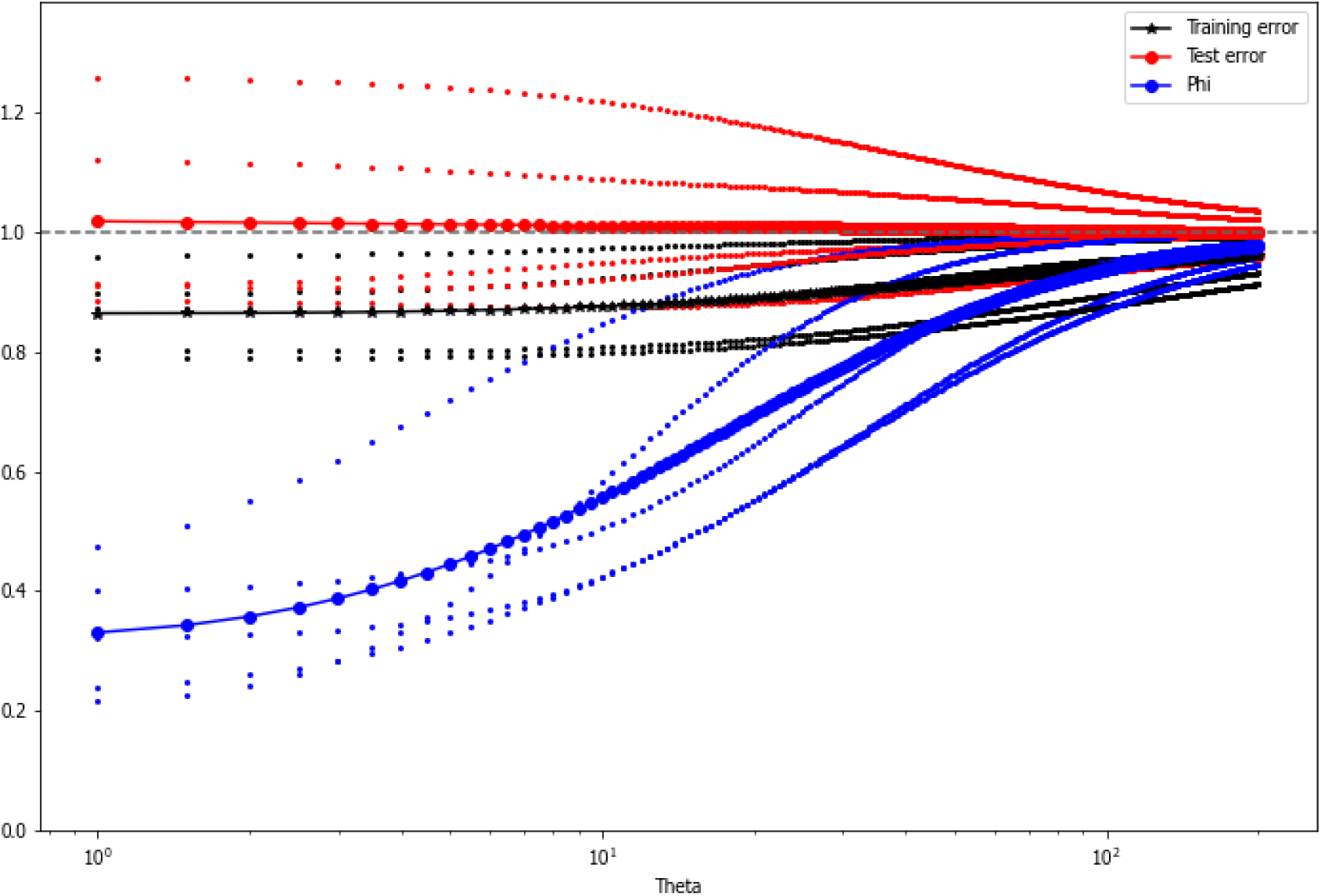
uCS-BME 5-fold cross-validation results that was used to choose the value of *θ* for PDB ID: 5KH8. The training and testing errors are shown in black and red, respectively. Shown in blue is *ϕ*, the effective ensemble size. The optimal *θ* is chosen as the value that minimizes the average testing error.

**Figure S11:**
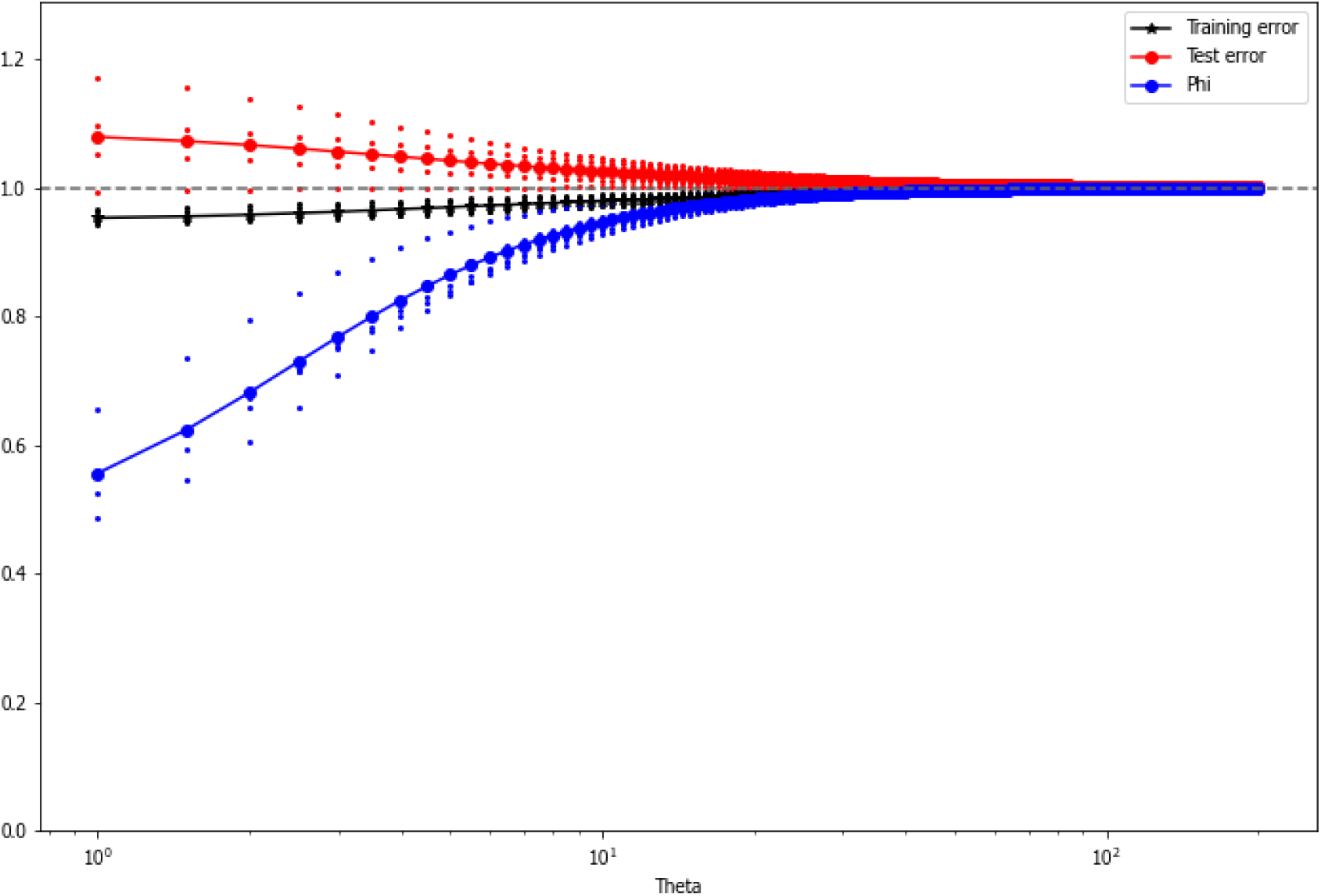
uCS-BME 5-fold cross-validation results that was used to choose the value of *θ* for PDB ID: 5KMZ. The training and testing errors are shown in black and red, respectively. Shown in blue is *ϕ*, the effective ensemble size. The optimal *θ* is chosen as the value that minimizes the average testing error.

**Figure S12:**
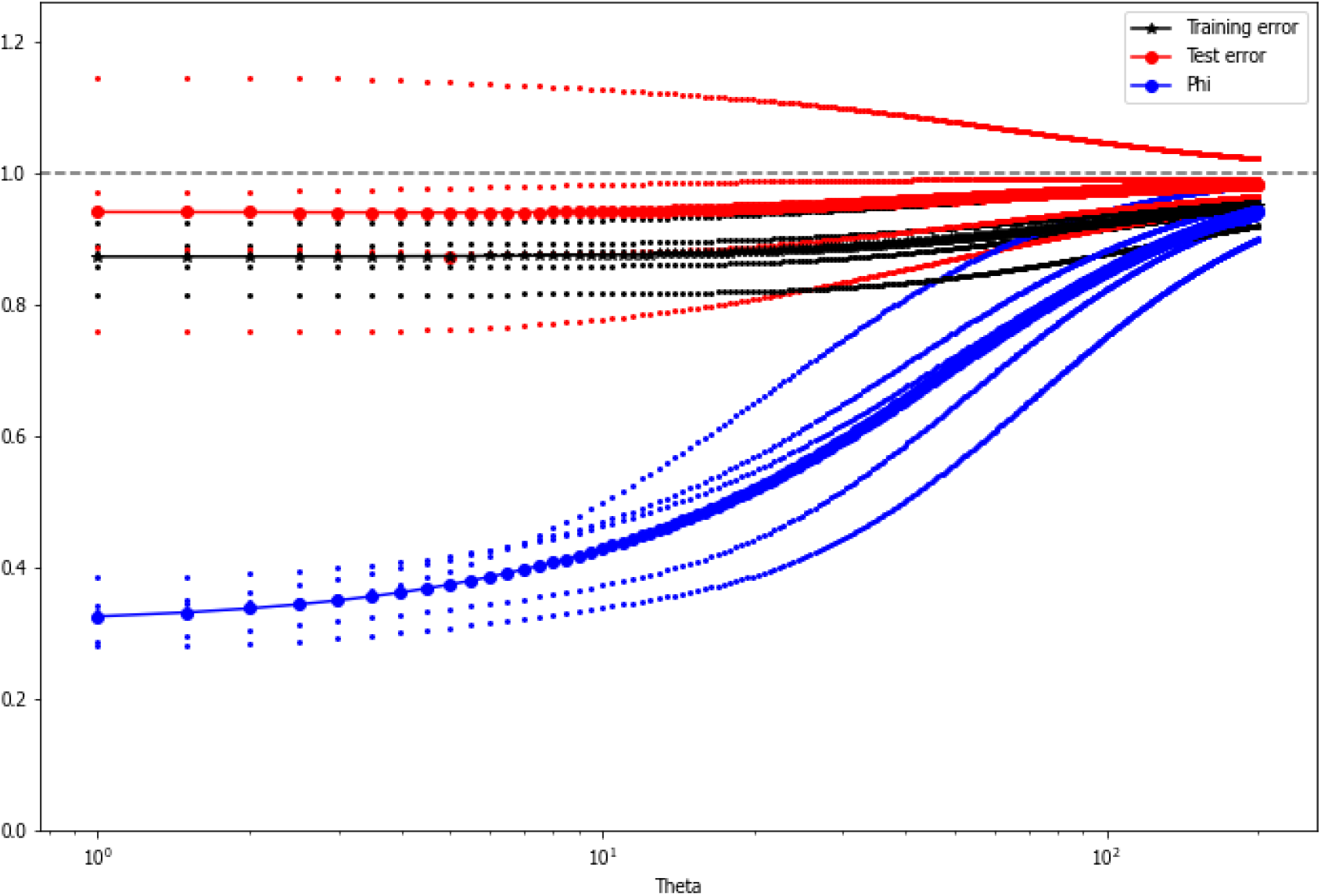
uCS-BME 5-fold cross-validation results that was used to choose the value of *θ* for PDB ID: 6GZK. The training and testing errors are shown in black and red, respectively. Shown in blue is *ϕ*, the effective ensemble size. The optimal *θ* is chosen as the value that minimizes the average testing error.

**Figure S13:**
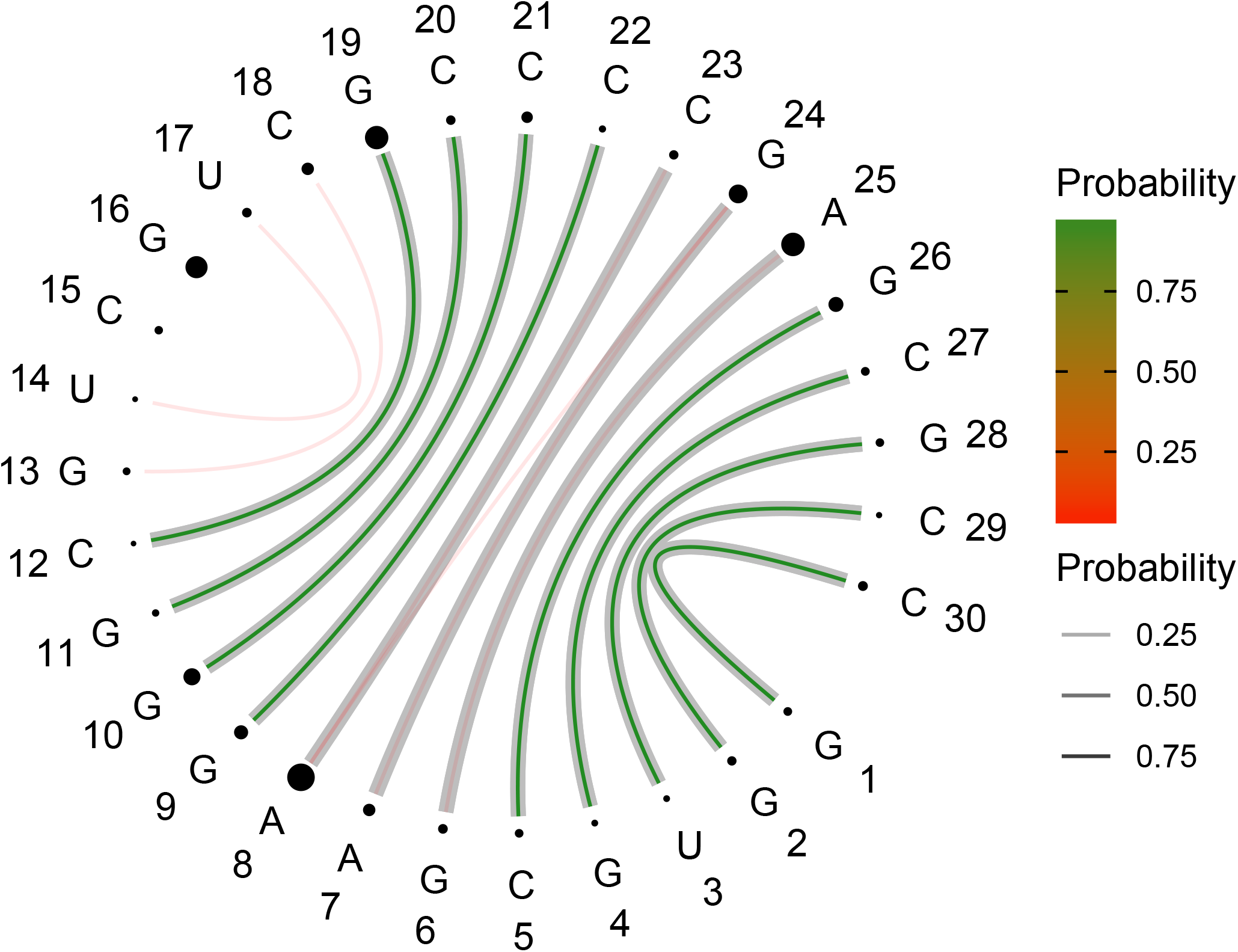
uCS-BME-derived Circular Secondary Structure Base Pairs Probabilities Plot (CS^2^BP^2^Plot) for PDB ID: 1HWQ. Base pair probabilities are color-coded; green-to-red associated with high-to-low base-pair probabilities. Probabilities are also encoded in the width of lines representing base pairs; the thicker the lines, the higher the base-pair probability. The thick lines that are shaded gray indicate base-pairs in the reference NMR model. The size of the spheres encode information about the relative residue-wise chemical shift error; the larger the sphere, the larger the chemical shift error associated with a given residue. Blue sphere indicate instances were no chemical shift data were able for the associated residue.

**Figure S14:**
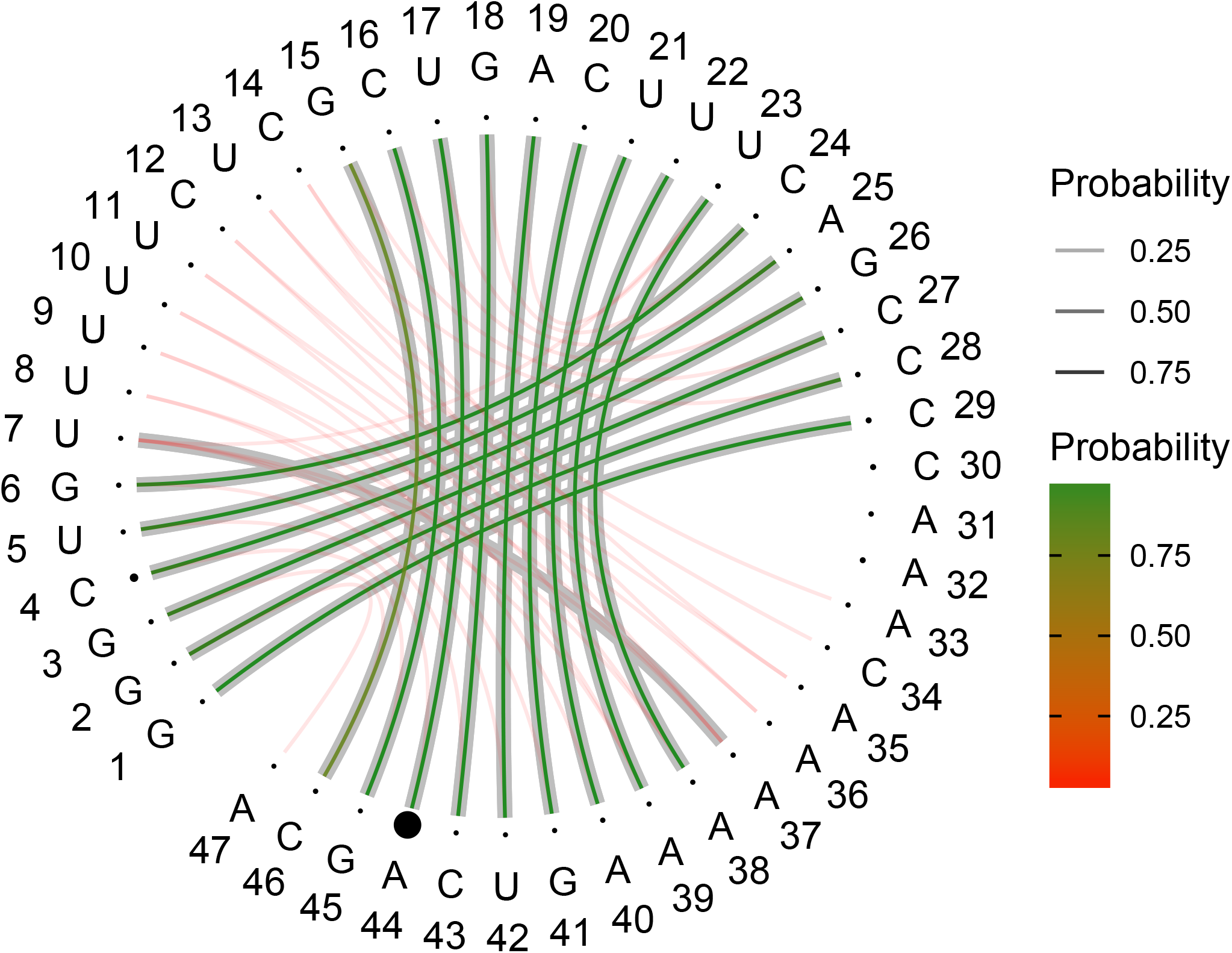
uCS-BME-derived Circular Secondary Structure Base Pairs Probabilities Plot (CS^2^BP^2^Plot) for PDB ID: 1YMO. Base pair probabilities are color-coded; green-to-red associated with high-to-low base-pair probabilities. Probabilities are also encoded in the width of lines representing base pairs; the thicker the lines, the higher the base-pair probability. The thick lines that are shaded gray indicate base-pairs in the reference NMR model. The size of the spheres encode information about the relative residue-wise chemical shift error; the larger the sphere, the larger the chemical shift error associated with a given residue. Blue sphere indicate instances were no chemical shift data were able for the associated residue.

**Figure S15:**
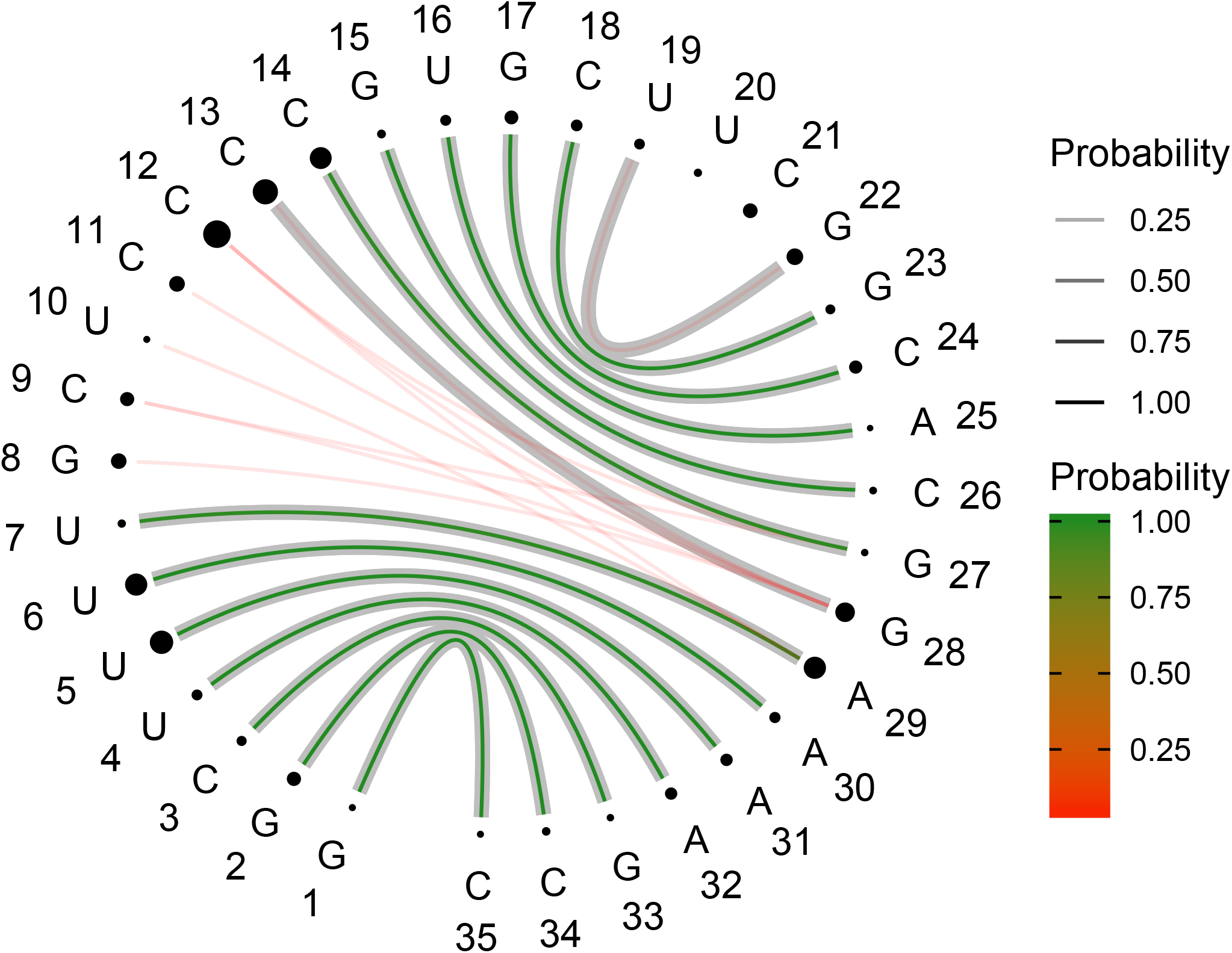
uCS-BME-derived Circular Secondary Structure Base Pairs Probabilities Plot (CS^2^BP^2^Plot) for PDB ID: 2L3E. Base pair probabilities are color-coded; green-to-red associated with high-to-low base-pair probabilities. Probabilities are also encoded in the width of lines representing base pairs; the thicker the lines, the higher the base-pair probability. The thick lines that are shaded gray indicate base-pairs in the reference NMR model. The size of the spheres encode information about the relative residue-wise chemical shift error; the larger the sphere, the larger the chemical shift error associated with a given residue. Blue sphere indicate instances were no chemical shift data were able for the associated residue.

**Figure S16:**
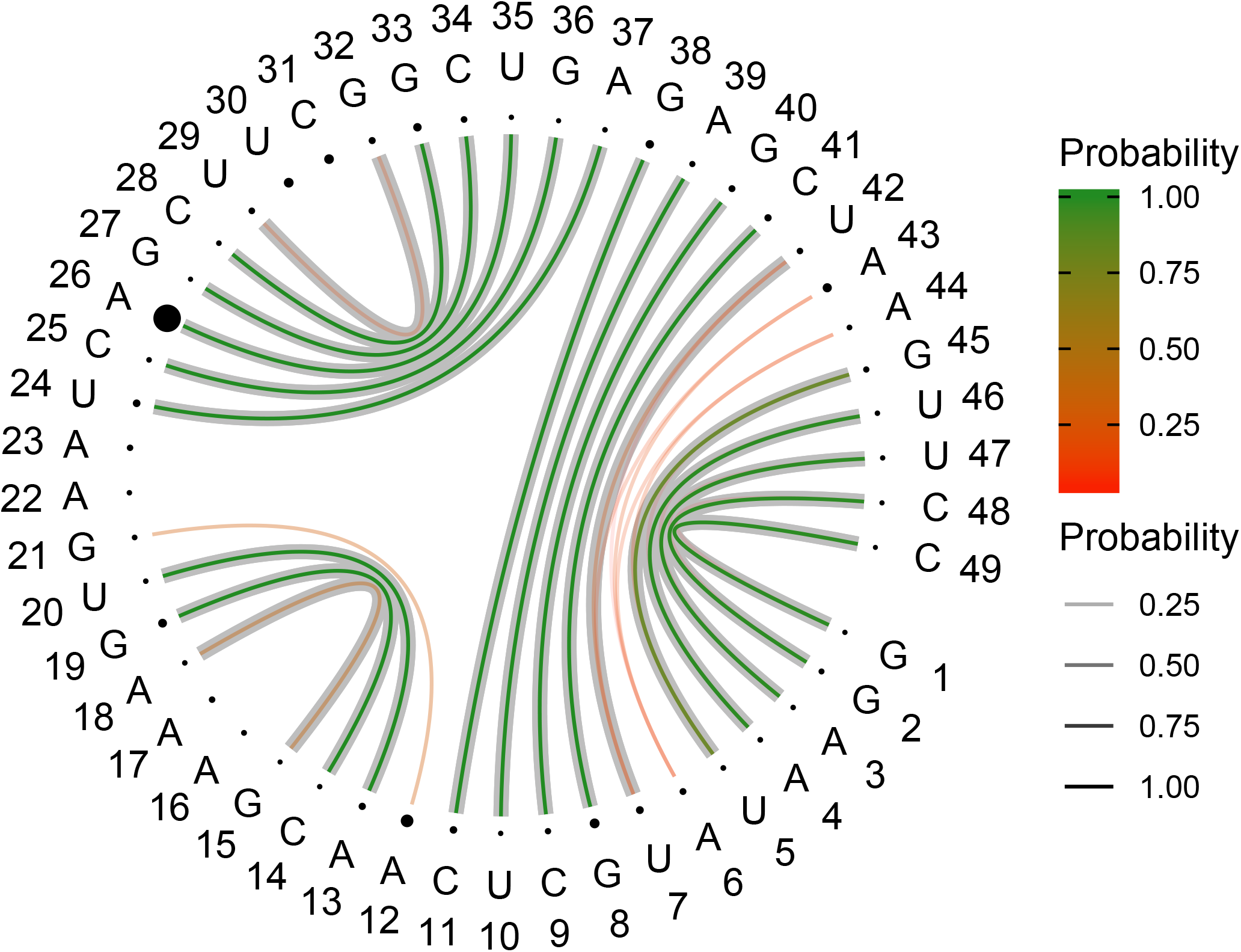
uCS-BME-derived Circular Secondary Structure Base Pairs Probabilities Plot (CS^2^BP^2^Plot) for PDB ID: 2LU0. Base pair probabilities are color-coded; green-to-red associated with high-to-low base-pair probabilities. Probabilities are also encoded in the width of lines representing base pairs; the thicker the lines, the higher the base-pair probability. The thick lines that are shaded gray indicate base-pairs in the reference NMR model. The size of the spheres encode information about the relative residue-wise chemical shift error; the larger the sphere, the larger the chemical shift error associated with a given residue. Blue sphere indicate instances were no chemical shift data were able for the associated residue.

**Figure S17:**
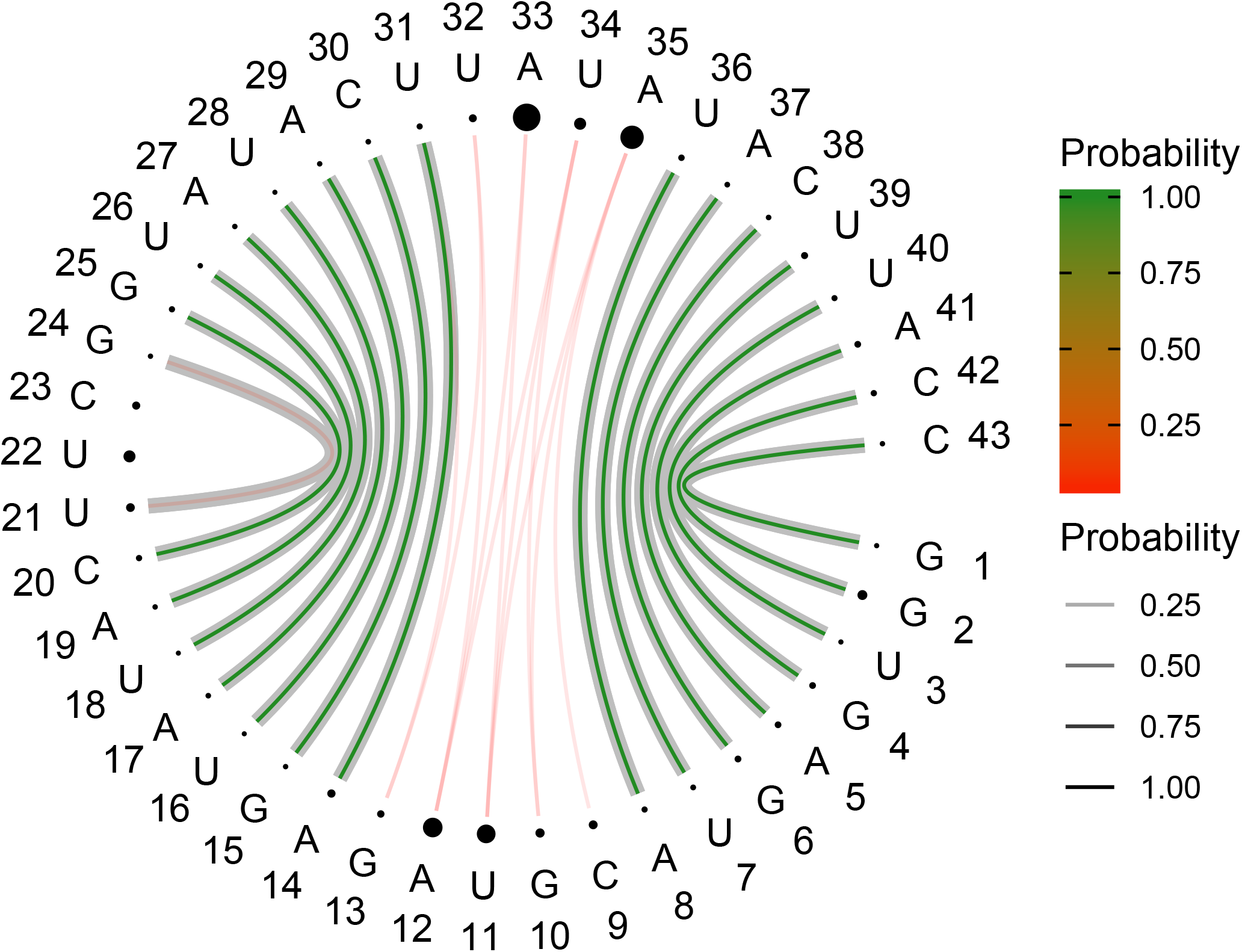
uCS-BME-derived Circular Secondary Structure Base Pairs Probabilities Plot (CS^2^BP^2^Plot) for PDB ID: 2N6X. Base pair probabilities are color-coded; green-to-red associated with high-to-low base-pair probabilities. Probabilities are also encoded in the width of lines representing base pairs; the thicker the lines, the higher the base-pair probability. The thick lines that are shaded gray indicate base-pairs in the reference NMR model. The size of the spheres encode information about the relative residue-wise chemical shift error; the larger the sphere, the larger the chemical shift error associated with a given residue. Blue sphere indicate instances were no chemical shift data were able for the associated residue.

**Figure S18:**
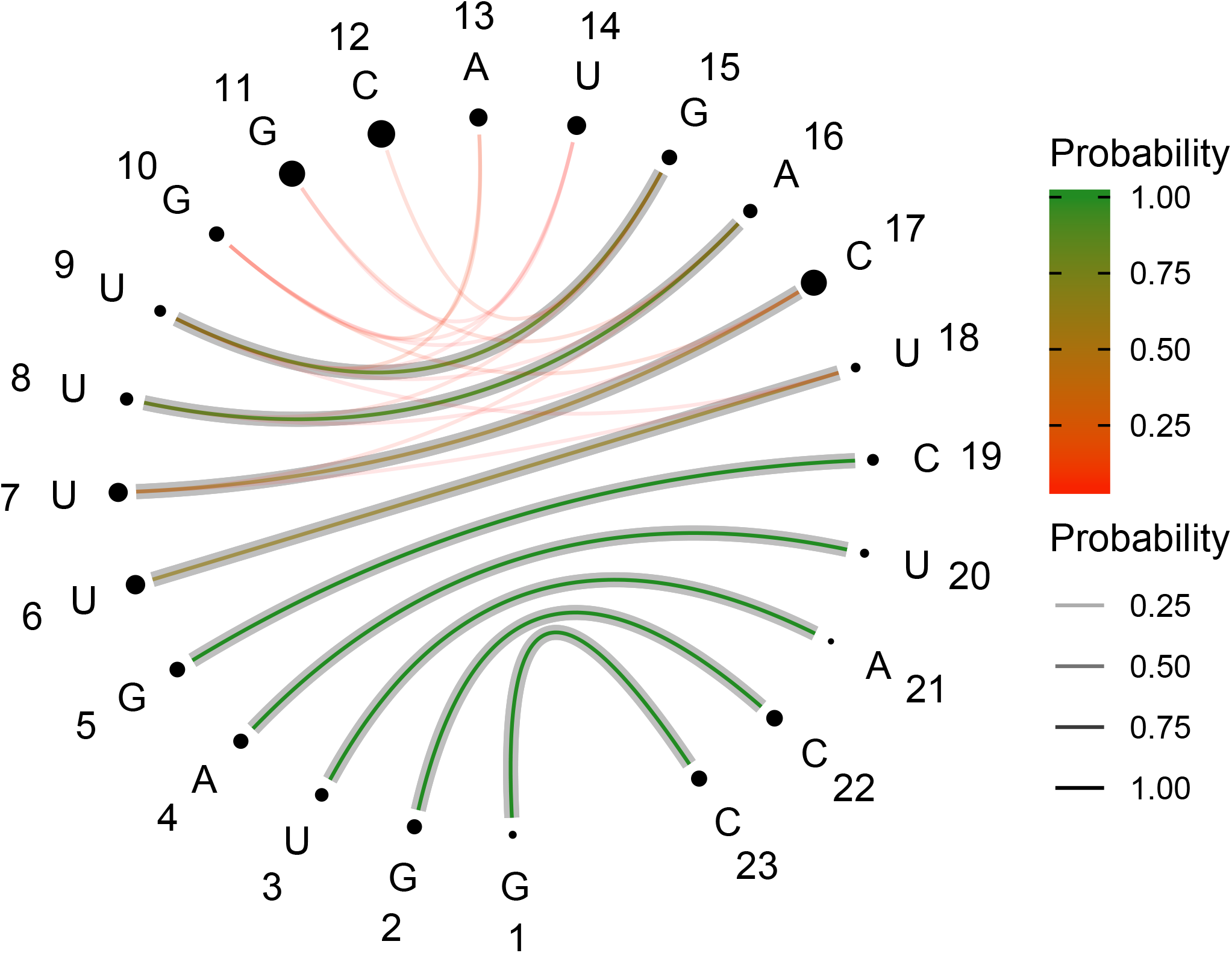
uCS-BME-derived Circular Secondary Structure Base Pairs Probabilities Plot (CS^2^BP^2^Plot) for PDB ID: 2N7X. Base pair probabilities are color-coded; green-to-red associated with high-to-low base-pair probabilities. Probabilities are also encoded in the width of lines representing base pairs; the thicker the lines, the higher the base-pair probability. The thick lines that are shaded gray indicate base-pairs in the reference NMR model. The size of the spheres encode information about the relative residue-wise chemical shift error; the larger the sphere, the larger the chemical shift error associated with a given residue. Blue sphere indicate instances were no chemical shift data were able for the associated residue.

**Figure S19:**
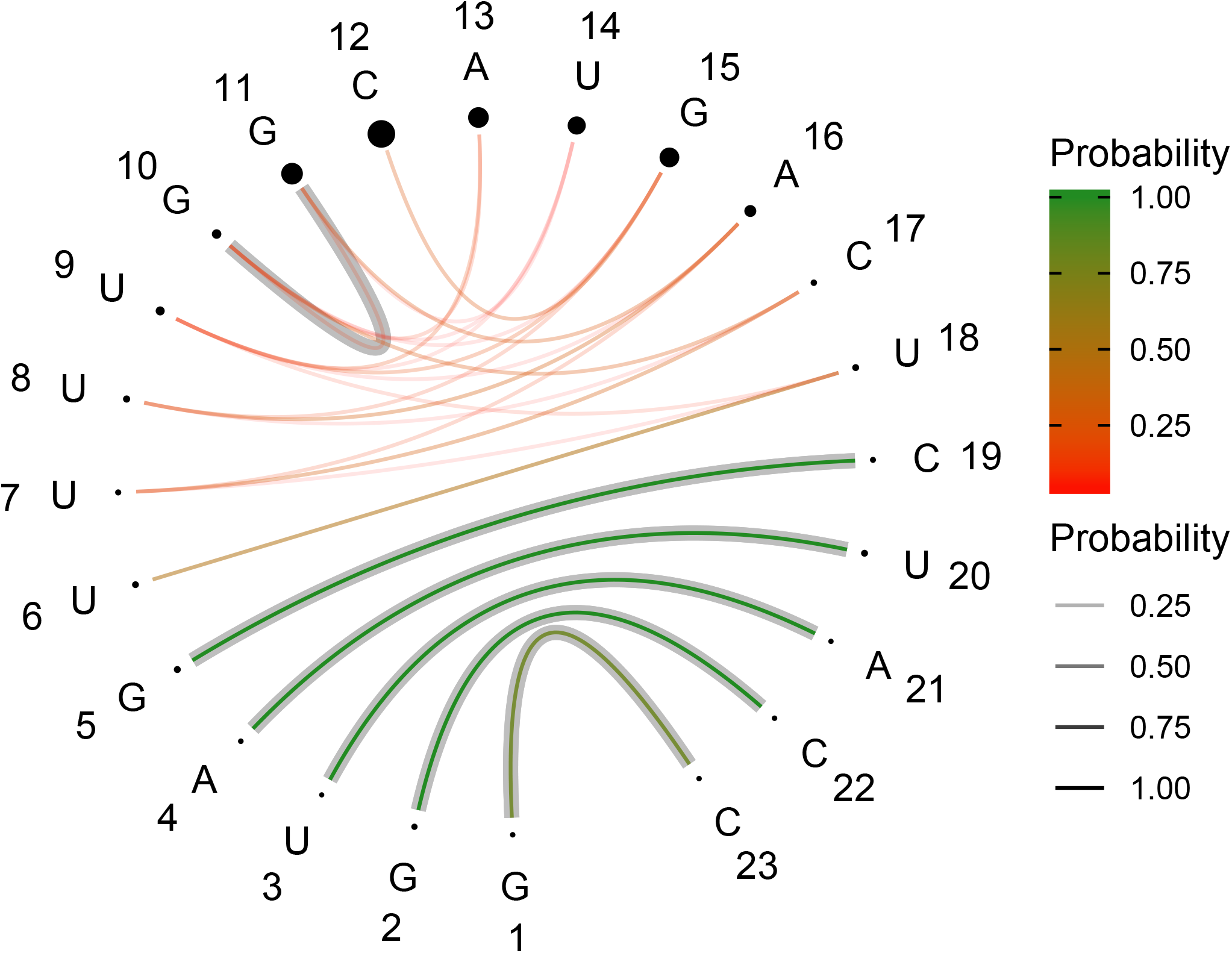
uCS-BME-derived Circular Secondary Structure Base Pairs Probabilities Plot (CS^2^BP^2^Plot) for PDB ID: 2N82. Base pair probabilities are color-coded; green-to-red associated with high-to-low base-pair probabilities. Probabilities are also encoded in the width of lines representing base pairs; the thicker the lines, the higher the base-pair probability. The thick lines that are shaded gray indicate base-pairs in the reference NMR model. The size of the spheres encode information about the relative residue-wise chemical shift error; the larger the sphere, the larger the chemical shift error associated with a given residue. Blue sphere indicate instances were no chemical shift data were able for the associated residue.

**Figure S20:**
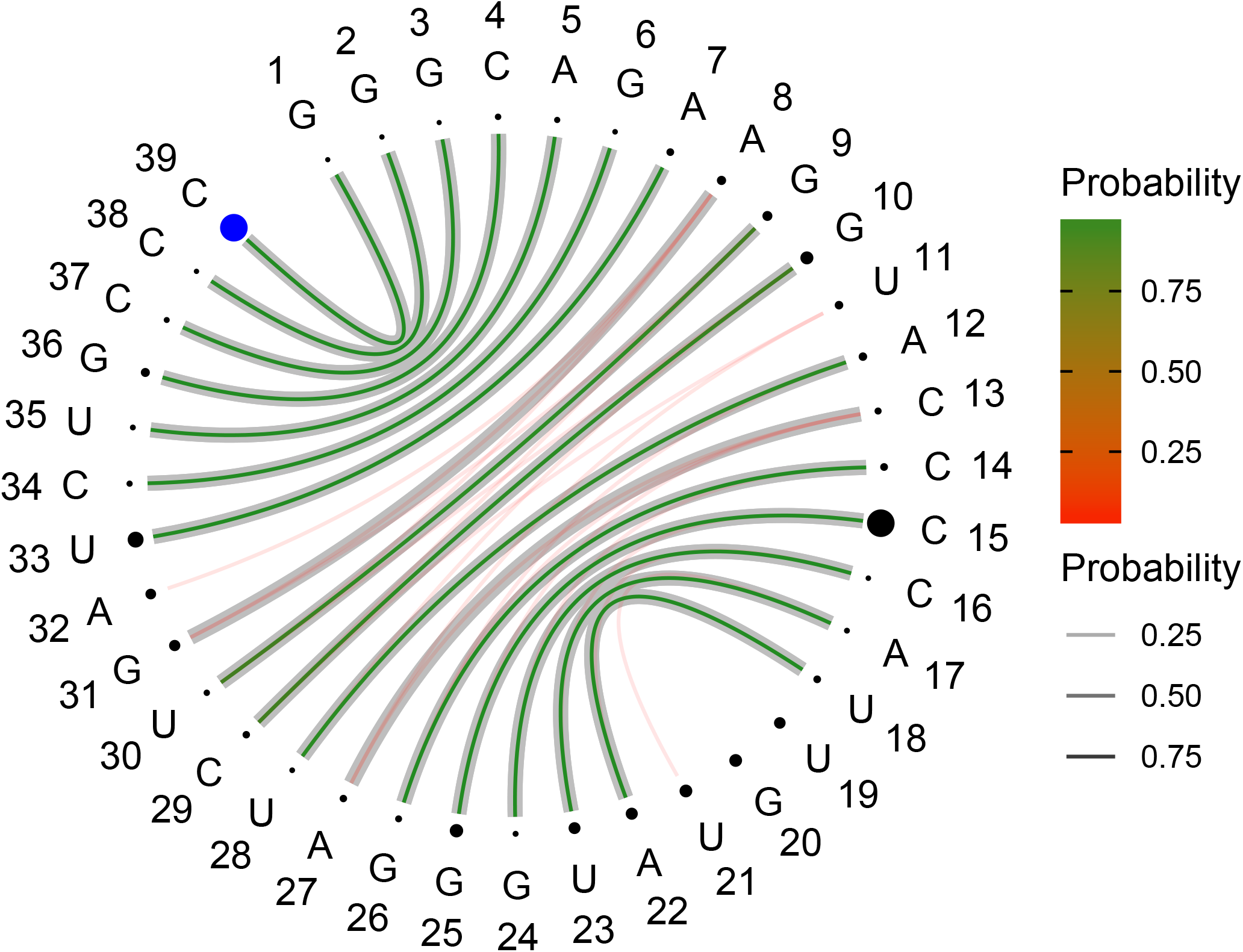
uCS-BME-derived Circular Secondary Structure Base Pairs Probabilities Plot (CS^2^BP^2^Plot) for PDB ID: 2NBY. Base pair probabilities are color-coded; green-to-red associated with high-to-low base-pair probabilities. Probabilities are also encoded in the width of lines representing base pairs; the thicker the lines, the higher the base-pair probability. The thick lines that are shaded gray indicate base-pairs in the reference NMR model. The size of the spheres encode information about the relative residue-wise chemical shift error; the larger the sphere, the larger the chemical shift error associated with a given residue. Blue sphere indicate instances were no chemical shift data were able for the associated residue.

**Figure S21:**
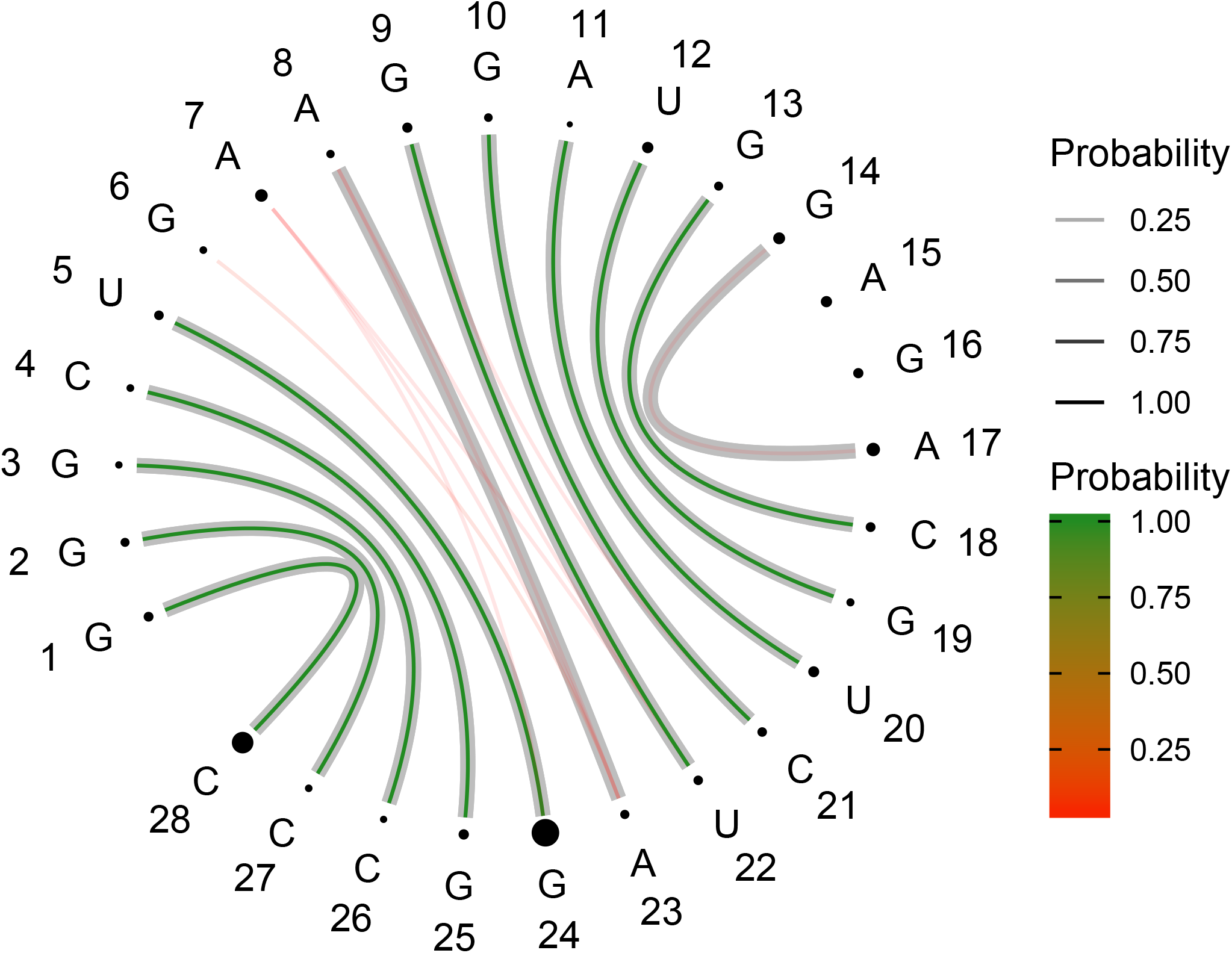
uCS-BME-derived Circular Secondary Structure Base Pairs Probabilities Plot (CS^2^BP^2^Plot) for PDB ID: 2NC0. Base pair probabilities are color-coded; green-to-red associated with high-to-low base-pair probabilities. Probabilities are also encoded in the width of lines representing base pairs; the thicker the lines, the higher the base-pair probability. The thick lines that are shaded gray indicate base-pairs in the reference NMR model. The size of the spheres encode information about the relative residue-wise chemical shift error; the larger the sphere, the larger the chemical shift error associated with a given residue. Blue sphere indicate instances were no chemical shift data were able for the associated residue.

**Figure S22:**
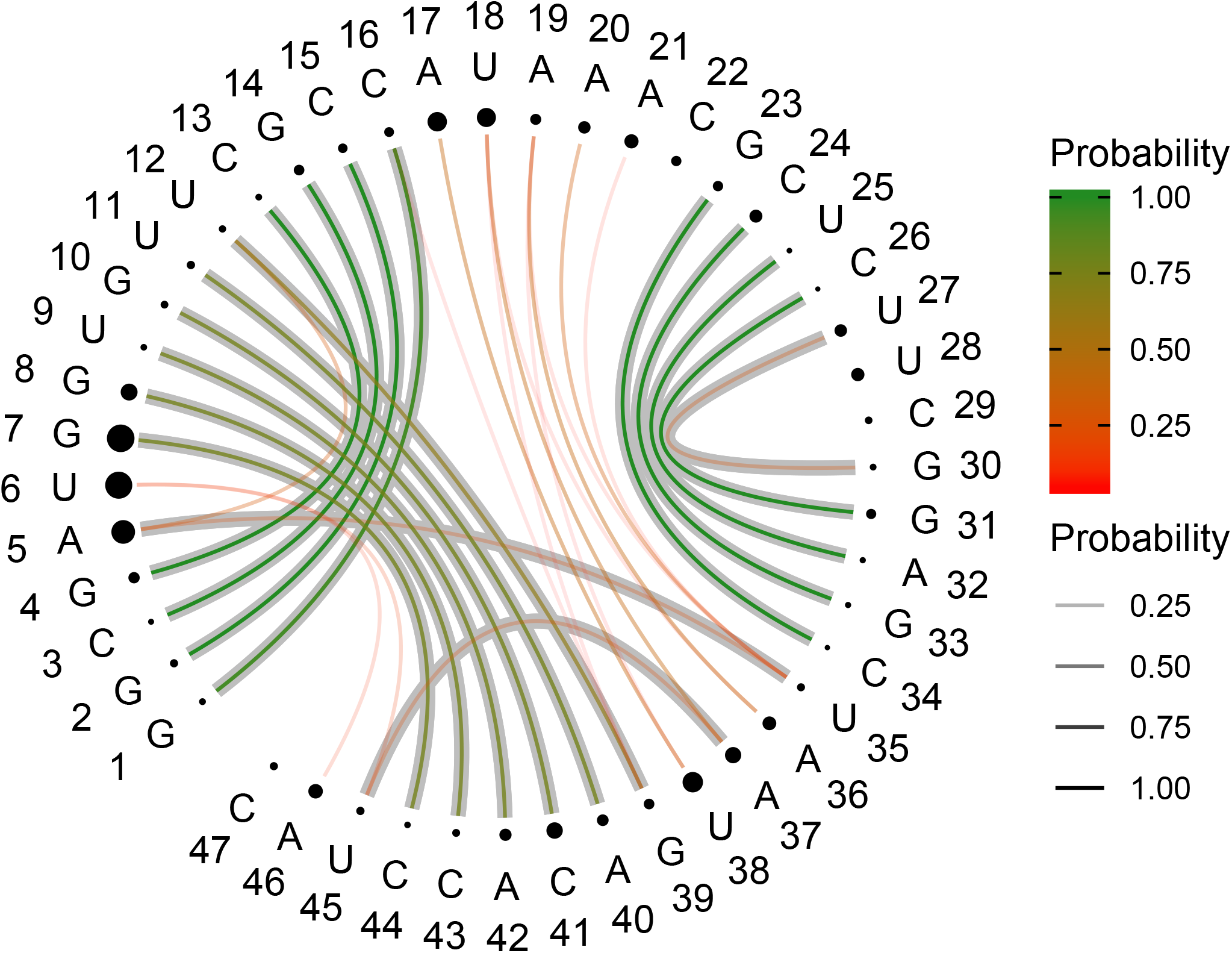
uCS-BME-derived Circular Secondary Structure Base Pairs Probabilities Plot (CS^2^BP^2^Plot) for PDB ID: 5KH8. Base pair probabilities are color-coded; green-to-red associated with high-to-low base-pair probabilities. Probabilities are also encoded in the width of lines representing base pairs; the thicker the lines, the higher the base-pair probability. The thick lines that are shaded gray indicate base-pairs in the reference NMR model. The size of the spheres encode information about the relative residue-wise chemical shift error; the larger the sphere, the larger the chemical shift error associated with a given residue. Blue sphere indicate instances were no chemical shift data were able for the associated residue.

**Figure S23:**
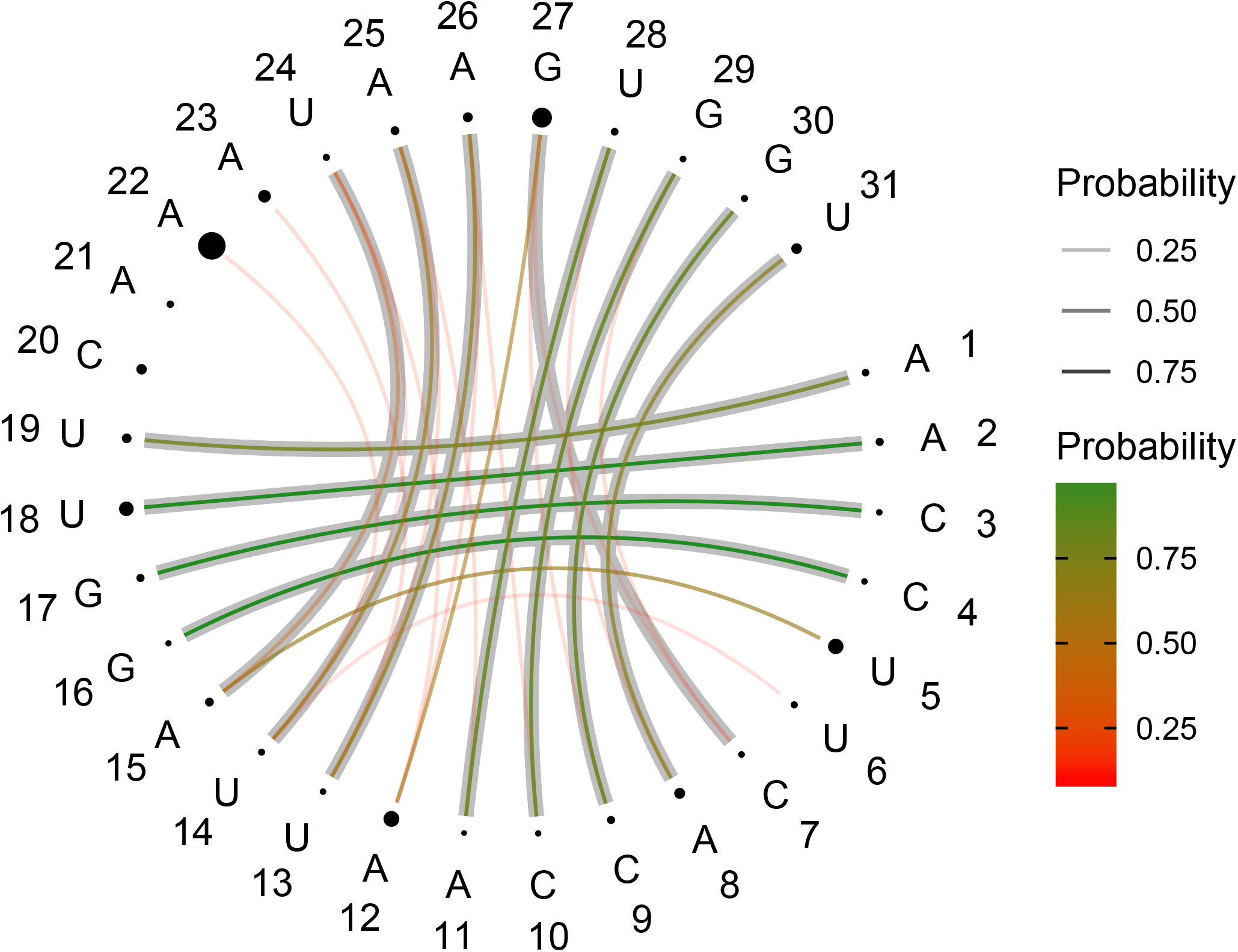
uCS-BME-derived Circular Secondary Structure Base Pairs Probabilities Plot (CS^2^BP^2^Plot) for PDB ID: 5KMZ. Base pair probabilities are color-coded; green-to-red associated with high-to-low base-pair probabilities. Probabilities are also encoded in the width of lines representing base pairs; the thicker the lines, the higher the base-pair probability. The thick lines that are shaded gray indicate base-pairs in the reference NMR model. The size of the spheres encode information about the relative residue-wise chemical shift error; the larger the sphere, the larger the chemical shift error associated with a given residue. Blue sphere indicate instances were no chemical shift data were able for the associated residue.

**Figure S24:**
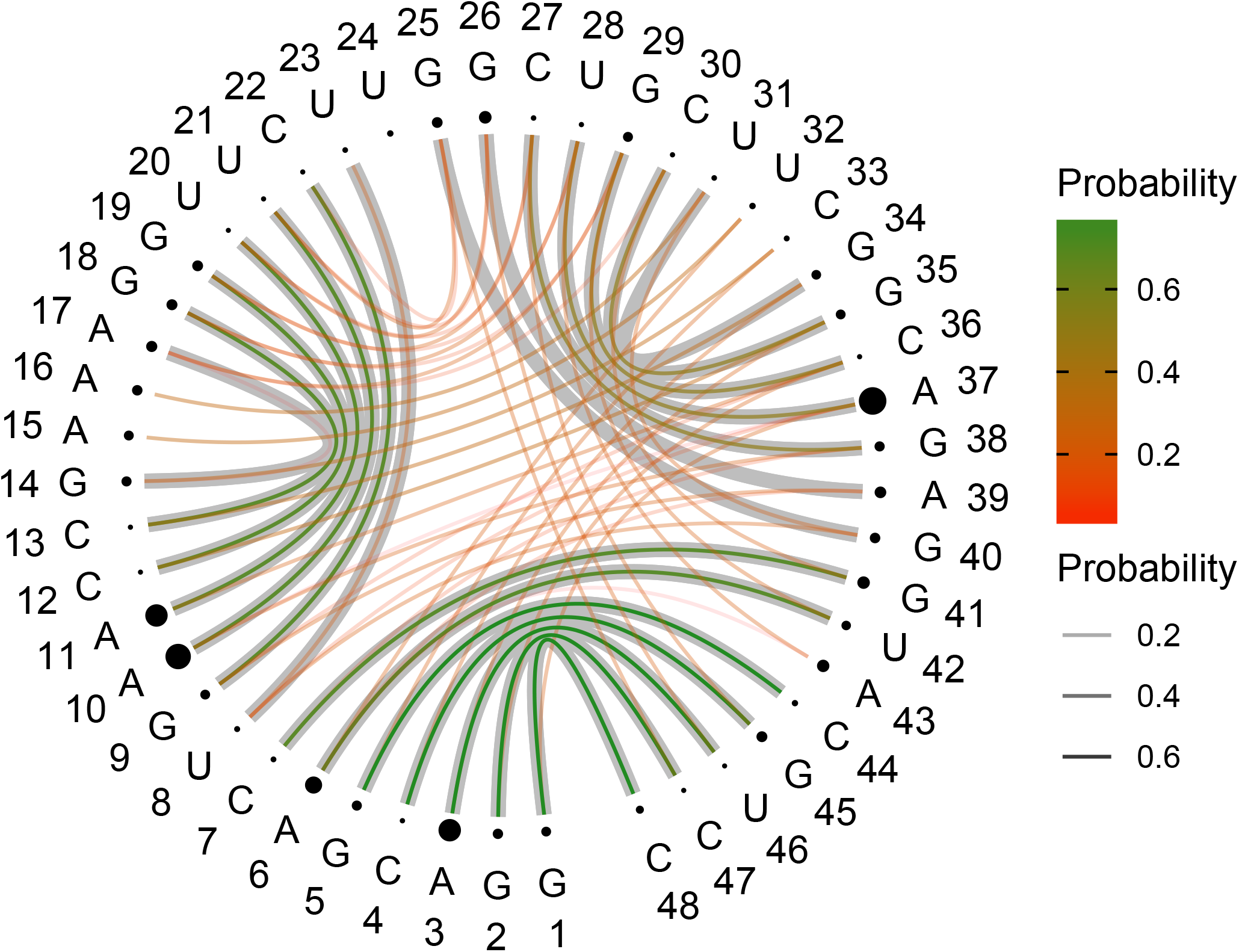
uCS-BME-derived Circular Secondary Structure Base Pairs Probabilities Plot (CS^2^BP^2^Plot) for PDB ID: 6GZK. Base pair probabilities are color-coded; green-to-red associated with high-to-low base-pair probabilities. Probabilities are also encoded in the width of lines representing base pairs; the thicker the lines, the higher the base-pair probability. The thick lines that are shaded gray indicate base-pairs in the reference NMR model. The size of the spheres encode information about the relative residue-wise chemical shift error; the larger the sphere, the larger the chemical shift error associated with a given residue. Blue sphere indicate instances were no chemical shift data were able for the associated residue.

**Figure S25:**
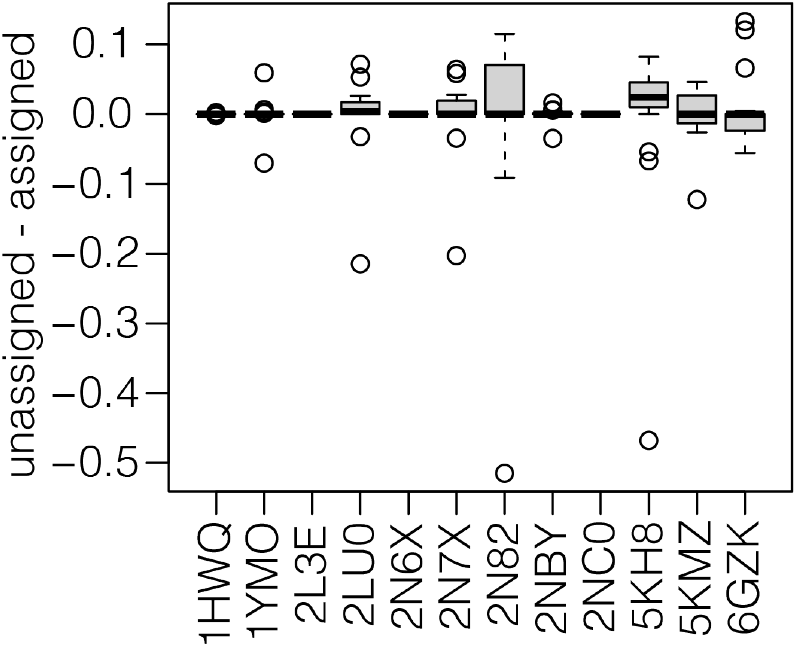
Boxplots of the differences between weights derived from CS-BME (assigned) and uCS-BME (unassigned).

**Table S1:** Estimated accuracy for SS2CS predictors trained to predict N1/H1 and N3/H3 chemical shift from structure (https://github.com/atfrank/SS2CS).

| N type | N<sub>N</sub> | MAE<sub>N</sub> (ppm) | H type | N<sub>H</sub> | MAE<sub>H</sub> (ppm) |
| --- | --- | --- | --- | --- | --- |
| N1 | 750 | 2.58 | H1 | 516 | 0.26 |
| N3 | 467 | 1.19 | H3 | 367 | 0.52 |
| mean | — | 1.89 | — | — | 0.39 |

**Table S2:** uCS-BME TPR for the SARS-CoV-2 3’-UTR as a function of the ^1^H (column) and ^15^N (row) bin size used to generate 2D histograms.

|  | 0.36 ppm | 0.37 ppm | 0.38 ppm | 0.39 ppm | 0.40 ppm | 0.41 ppm | 0.42 ppm |
| --- | --- | --- | --- | --- | --- | --- | --- |
| 1.86 ppm | 77.71 | 77.71 | 77.07 | 77.07 | 77.07 | 77.07 | 78.21 |
| 1.87 ppm | 77.71 | 77.71 | 77.07 | 77.07 | 77.07 | 77.07 | 78.21 |
| 1.88 ppm | 78.21 | 77.71 | 77.07 | 77.07 | 77.07 | 77.07 | 78.21 |
| 1.89 ppm | 77.71 | 77.71 | 77.07 | 77.07 | 77.07 | 77.07 | 78.21 |
| 1.90 ppm | 77.71 | 77.71 | 77.07 | 77.07 | 77.07 | 77.07 | 78.21 |
| 1.91 ppm | 77.71 | 77.71 | 77.07 | 77.07 | 77.07 | 77.07 | 78.21 |
| 1.92 ppm | 77.71 | 77.71 | 77.07 | 77.07 | 77.07 | 77.07 | 78.21 |

**Table S3:** uCS-BME PPV for the SARS-CoV-2 3’-UTR as a function of the ^1^H (column) and ^15^N (row) bin size used to generate 2D histograms.

|  | 0.36 ppm | 0.37 ppm | 0.38 ppm | 0.39 ppm | 0.40 ppm | 0.41 ppm | 0.42 ppm |
| --- | --- | --- | --- | --- | --- | --- | --- |
| 1.86 ppm | 73.74 | 92.08 | 92.08 | 92.08 | 93.07 | 93.07 | 93.07 |
| 1.87 ppm | 74.26 | 92.08 | 92.08 | 92.08 | 93.07 | 93.07 | 93.07 |
| 1.88 ppm | 74.26 | 92.08 | 92.08 | 92.08 | 93.07 | 93.07 | 93.07 |
| 1.89 ppm | 74.26 | 92.08 | 92.08 | 92.08 | 93.07 | 93.07 | 93.07 |
| 1.90 ppm | 74.26 | 92.08 | 92.08 | 92.08 | 93.07 | 93.07 | 93.07 |
| 1.91 ppm | 74.26 | 92.08 | 92.08 | 92.08 | 93.07 | 93.07 | 93.07 |
| 1.92 ppm | 74.26 | 92.08 | 92.08 | 92.08 | 93.07 | 93.07 | 93.07 |

**Table S4:** uCS-BME TPR for the SARS-CoV-2 5’-UTR as a function of the ^1^H (column) and ^15^N (row) bin size used to generate 2D histograms.

|  | 0.36 ppm | 0.37 ppm | 0.38 ppm | 0.39 ppm | 0.40 ppm | 0.41 ppm | 0.42 ppm |
| --- | --- | --- | --- | --- | --- | --- | --- |
| 1.86 ppm | 88.41 | 88.41 | 87.68 | 87.68 | 87.68 | 87.68 | 88.41 |
| 1.87 ppm | 88.41 | 88.41 | 87.68 | 87.68 | 87.68 | 87.68 | 88.41 |
| 1.88 ppm | 88.41 | 88.41 | 87.68 | 87.68 | 87.68 | 87.68 | 88.41 |
| 1.89 ppm | 88.41 | 88.41 | 87.68 | 87.68 | 87.68 | 87.68 | 88.41 |
| 1.90 ppm | 88.41 | 88.41 | 87.68 | 87.68 | 87.68 | 87.68 | 88.41 |
| 1.91 ppm | 88.41 | 88.41 | 87.68 | 87.68 | 87.68 | 87.68 | 88.41 |
| 1.92 ppm | 88.41 | 88.41 | 87.68 | 87.68 | 87.68 | 87.68 | 88.41 |

**Table S5:** uCS-BME PPV for the SARS-CoV-2 5’-UTR as a function of the ^1^H (column) and ^15^N (row) bin size used to generate 2D histograms.

|  | 0.36 ppm | 0.37 ppm | 0.38 ppm | 0.39 ppm | 0.40 ppm | 0.41 ppm | 0.42 ppm |
| --- | --- | --- | --- | --- | --- | --- | --- |
| 1.86 ppm | 69.52 | 88.57 | 88.57 | 88.57 | 89.52 | 89.52 | 89.52 |
| 1.87 ppm | 71.43 | 88.57 | 88.57 | 88.57 | 89.52 | 89.52 | 89.52 |
| 1.88 ppm | 71.43 | 88.57 | 88.57 | 88.57 | 89.52 | 89.52 | 89.52 |
| 1.89 ppm | 71.43 | 88.57 | 88.57 | 88.57 | 89.52 | 89.52 | 89.52 |
| 1.90 ppm | 71.43 | 88.57 | 88.57 | 88.57 | 89.52 | 89.52 | 89.52 |
| 1.91 ppm | 71.43 | 88.57 | 88.57 | 88.57 | 89.52 | 89.52 | 89.52 |
| 1.92 ppm | 71.43 | 88.57 | 88.57 | 88.57 | 89.52 | 89.52 | 89.52 |

